# Integrated cardio-behavioural defensive states

**DOI:** 10.1101/2022.09.22.509009

**Authors:** Jérémy Signoret-Genest, Nina Schukraft, Sara L. Reis, Dennis Segebarth, Philip Tovote

**Affiliations:** Institute of Clinical Neurobiology, University Hospital Wuerzburg, Wuerzburg, Germany; Center for Mental Health; University Hospital Wuerzburg, Wuerzburg, Germany

## Abstract

Fear and anxiety are brain states that evolved to mediate defensive responses to threat. While it is clear that the defense reaction includes multiple interacting behavioural, autonomic and endocrine adjustments, their integrative nature is poorly understood. In particular, threat has been associated with various cardiac changes, yet a clear consensus on their relevance for the integrated defense reaction is missing. We here define rapid microstates associated with specific behaviours and heart rate dynamics, both affected by long-lasting macrostates and reflecting context-dependent threat levels. In addition, we demonstrate that one of the most commonly used defensive behavioural responses, freezing measured by immobility, is part of an integrated cardio-behavioural microstate mediated by specific midbrain circuit elements. Our work puts forth a framework for systematic integration of cardiac and behavioural readouts that presents the basis for a better understanding of complex neural defensive states and their associated systemic functions.

## Main

The defense reaction in response to threat, a central element of human fear or anxiety, encompasses multiple behavioural, autonomic and endocrine adjustments, controlled and integrated by neural circuits of nervous systems^1^. Based almost exclusively on behavioural responses, a unifying, across-species concept describes the dynamic nature of defensive responses as ‘states’^2–4^. However, by focusing on behavioural dynamics alone, a comprehensive understanding of defensive states and the integrated nature of their individual components remains incomplete. Yet, this is critical to causally link the known variety of neuronal states to their systemic readouts, such as behaviour^5^. Brain circuitry involving the midbrain periaqueductal grey (PAG) has long been suggested to play a major role in mediating defensive states by integrating behavioural and cardiac components^6–9^, but a lack of integrated analyses has prevented clarification of precise circuit mechanisms.

While investigations of defensive states predominantly focused on threat-induced behavioural changes^10, 11^, numerous studies throughout the years also addressed ‘autonomic’ adaptations to threats, in particular changes in heart rate (HR)^12–15^. Similar to behavioural responses, HR is considered the product of multiple internal processes sensitive to threat. In contrast to relatively robust defensive behaviours elicited under tightly controlled experimental conditions, studies on defensive autonomic responses have yielded complex, paradoxical observations and sometimes seemingly contradictory findings. Under threat conditions, both deceleration (bradycardia) as well acceleration (tachycardia) of HR have been reported^8, 13, 16–23^. Overall, the heterogeneous results have divided the field into research either ignoring cardiac readouts as quantifiable measures for defensive states or, on the other hand, equating specific HR responses directly with fear.

Based on novel analyses of a large dataset of concomitant behavioural, HR and thermal measures during various behavioural paradigms in freely moving mice, we here define transient *microstates* and their interaction with longer-lasting *macrostates* to explain tachycardic as well bradycardic defensive responses and associated behavioural patterns. In addition to revealing cue- and context-dependency of integrated defense states, HR indices precisely identify defensive state transitions as well as contextual threat levels. Furthermore, optogenetic perturbational approaches enable assignment of particular ‘state generator’ roles to individual neural circuit elements within the midbrain periaqueductal grey (PAG). We here introduce a novel framework for characterization of integrated cardio-behavioural defensive states that serves as the basis for a comprehensive understanding of complex neuronal mechanisms underlying aversive emotions such as fear and anxiety.

### Heart rate changes as determinants of defensive micro- and macrostates

As a prerequisite for understanding integrated defensive states, we used a recently developed conditioned flight paradigm in which a serial compound stimulus (SCS), an auditory cue (pure tone followed by white noise) is paired with a mild electrical footshock, eliciting rapid switches in behavioural state^24^ (Extended Data Fig. 1c). Freely moving mice implanted with ECG electrodes (Fig. 1a, Extended Data Fig. 1a,b) exhibited stimulus-specific combinations of HR and behavioural responses: the pure tone period was characterized by immobility and a decrease in HR, while white noise was accompanied by interspersed flight/immobility and an increase in HR (Fig. 1b). HR during spontaneous, non CS-evoked freezing episodes decreased likewise (Fig. 1c), suggesting the existence of a stereotypical transient cardio-behavioural *microstate* characterized by immobility and HR decrease, i.e. bradycardia. However, the amplitude of immobility-associated bradycardia increased progressively throughout the conditioning session (Fig. 1d,e).

**Fig. 1.**
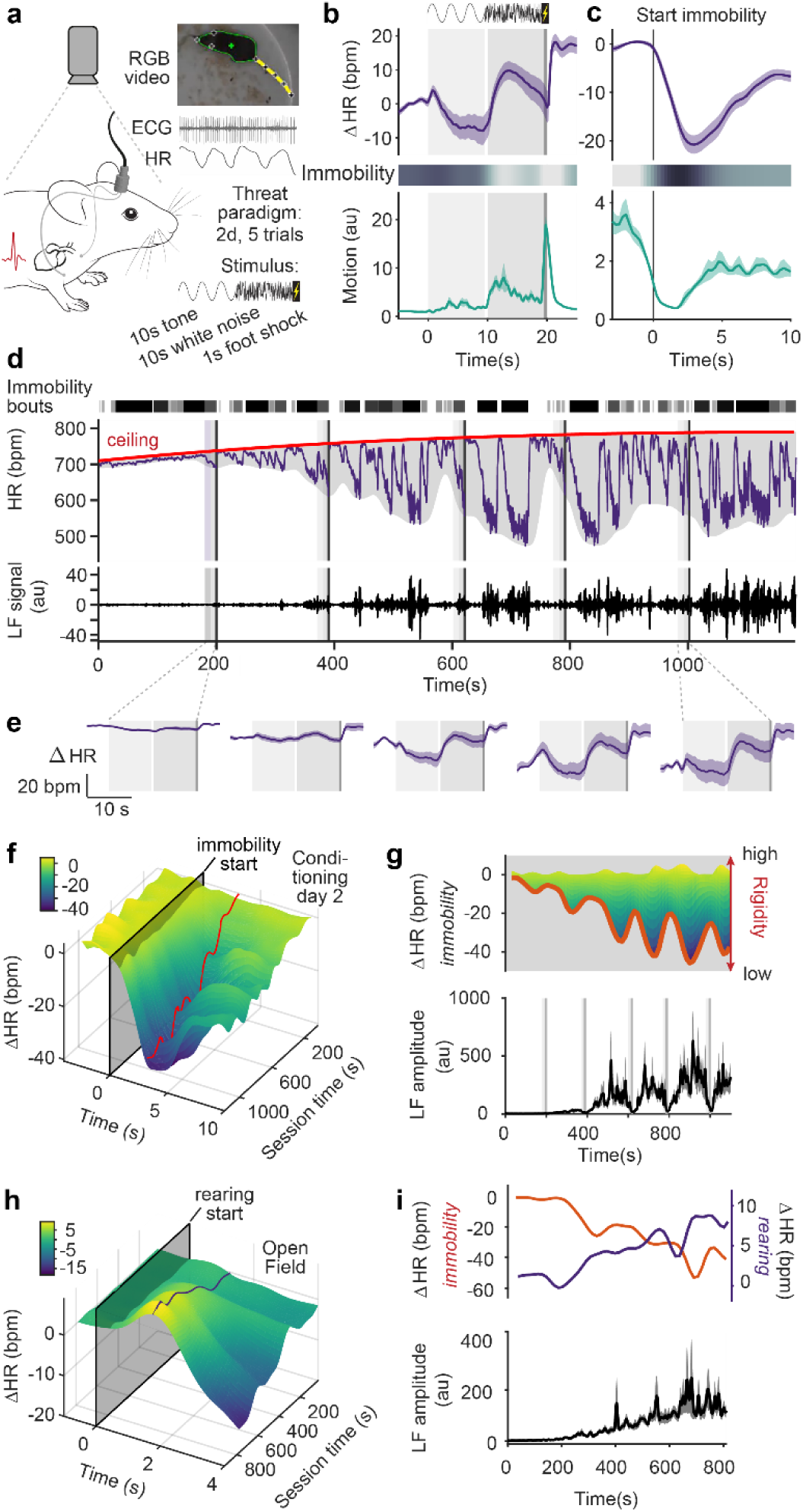
Heart rate changes reflect the interaction between defensive micro- and macrostates. **a**, Electrocardiograms were recorded from video-tracked freely moving mice to concomitantly study HR readouts and behaviour in a two-day fear conditioning paradigm for which a compound stimulus (tone then white noise) is paired with a shock. **b-c,** PSTHs showing the average HR (top) and behavioural responses (middle and bottom) to the 5 CS-US pairings during the second day of conditioning (**b**, n = 33 mice) or spontaneous immobility episodes (**c**, n = 33 mice); in both cases, immobility was associated with pronounced bradycardia. **d,** Example trace from a single mouse for a conditioning day 2 recording. Shaded rectangles indicate CS-US pairings (five in total). Raw HR (middle) illustrates how bradycardic events are clearly correlated with individual immobility bouts (dashed lines on top). It also shows that HR progressively increases from the beginning of the recording, following a latent maximum, the *ceiling* (red curve). The overall amplitude of the HR changes increases over time, as visualized by the smoothed envelope in grey. Finally, HR presents oscillations in the LF range, variability which is obvious during bradycardic episodes (middle curve), or after isolating the corresponding signal from the raw HR (bottom curve). **e,** PSTHs showing the average HR-to-ceiling traces for each of the five individual CS-US pairings (n = 33 mice), on which the progressive increase in amplitude is clearly visible. **f**, 3D representation showing the increase in amplitude of the immobility-associated HR changes in function of the recording time for conditioning day 2. The red line represents the lowest value for each time point (i.e. bradycardia’s peak value). **g,** top, side view from **f** to better show the dynamics. The amplitude of immobility-associated bradycardia over time (red curve, same as (**f**)) reflects *rigidity*: when *rigidity* is high (beginning of the recording), immobility-associated bradycardia is constrained to low amplitude. Conversely, progressive increase in that amplitude identifies lower *rigidity*. Bottom, average quantification of the LF amplitude (n = 33 mice), also developing over time and appearing to be correlated to *rigidity*. **h,** 3D representation showing the increase in rearing-associated HR changes along time in the OF (n = 23 mice). **i,** top, curves of immobility- and rearing-associated HR changes during OF recordings; bottom, LF amplitude during the same recordings (n = 23 mice), showing that rigidity similarly affects HR changes related to different behaviours and LF in different contexts.

How can we explain such a progressive change in the microstate level? Crucially, it affected the cardiac component of both spontaneous and CS-evoked immobility-bradycardia microstate (Fig. 1d-f), and was paralleled by low-frequency (LF) HR variability fluctuations, a murine equivalent to the LF band of human HR variability, which has been associated with emotional states^25^ (Fig. 1d, **bottom panel**). This led us to hypothesise that one or several underlying processes were interfering with HR dynamics at a more global level, and in particular with the *expression* of the cardiovascular component of the defensive microstate. Overall, HR evolved within the boundaries of a smooth envelope, where minimum values mainly corresponded to the HR decreases associated with immobility episodes, while maximum values never exceeded a monotonic curve (Fig. 1d, **top panel**). Because those changes operated at extended timescales when compared with short-lasting, e.g. cue-induced immobility bouts, we refer to them as dynamic *macrostates*. To capture the slowly shifting upper boundary of HR, we defined the *ceiling* macrostate, operationalized by latent maximum HR value, therefore progressively increasing in our conditions. In contrast, we attribute changes of the lower HR boundary to the *rigidity* macrostate, which constrains the range of values that HR can reach by opposing changes. This is reflected by the initially small amplitude of LF oscillations and immobility-associated bradycardia, which increases over time (Fig. 1e-g). Important to note, while *ceiling* increased, *rigidity* decreased across the session.

Beyond threat conditioning, we wondered if our macrostates were 1) also present in other contexts, and therefore dependent on general underlying processes and 2) affected other microstates in a similar fashion. Using the commonly used Open Field Test (OF), we found another microstate, in which rearing was associated with a transient increase in HR (Fig. 1h and Extended Data Fig. 1d). The amplitude of that tachycardia increased over the course of the recording and was correlated with amplitude increases in LF as well as immobility-associated bradycardia (Fig. 1i), similar to the progressive changes in immobility-associated bradycardia during conditioning. This confirms that the *rigidity* macrostate reflects a general mechanism affecting cardiac responses under threat conditions.

Taken together, these findings suggest that HR dynamics do not simply follow changes in behavioural activity, but reflect integrated defensive microstates with behavioural and autonomic components. Furthermore, we identified slow macrostate changes that exert strong influences on moment-to-moment microstates such as those induced by threatening cues.

### Defensive state dynamics determine cardiac responses

A corollary to our finding that the *rigidity* macrostate affects e.g. cue-induced defensive microstates, is that quantifying the latter without taking the former into account leads to confounded results and added variability. To circumvent this issue of added complexity by interaction of micro- and macrostates, other studies have for a long time introduced handling procedures^26^ or ‘acclimation’ periods before recordings, thereby attempting to remove influence of macrostates experimentally. However, in parallel to evoking repeated and poorly controlled stressful events beforehand, this means essentially manipulating the defensive state itself, and ultimately ignoring potentially important aspects of defensiveness. On the other hand, the slow increase in “baseline” HR level (*ceiling*) could confound quantifications based on absolute values. The commonly used moment-to-moment HR differences or changes from a baseline (delta HR) remove the influence of macrostates analytically at the cost of losing information about absolute HR. We therefore introduced a detrended ‘HR-to-ceiling’ measure (delta of HR at time t and the ceiling) as a representation of HR free from the influence of the *ceiling* macrostate and inter-mice differences in basal HR, but allowing for direct comparisons between conditions and/or individuals (Fig. 2a). Strikingly, LF amplitude correlated more strongly with HR-to-ceiling values than absolute HR (Fig. 2b and Extended Data Fig. 2a). LF oscillations, which correspond to a form of HR variability (HRV), are thought to reflect complex interactions between the parasympathetic and sympathetic systems^27–29^. The fact that HR-to-ceiling correlates to LF better than absolute HR therefore suggests that it has not only analytical but also biological relevance. In contrast to what delta HR suggested, this approach revealed that bradycardia is present with an extremely low amplitude also during immobility bouts early in the session (SCS1) and that it also occurs concomitantly with immobility during late tone exposures (SCS5), thereby confirming the global progressive increase in amplitude (Fig. 2c, Extended Data Fig. 2c-e).

**Fig. 2.**
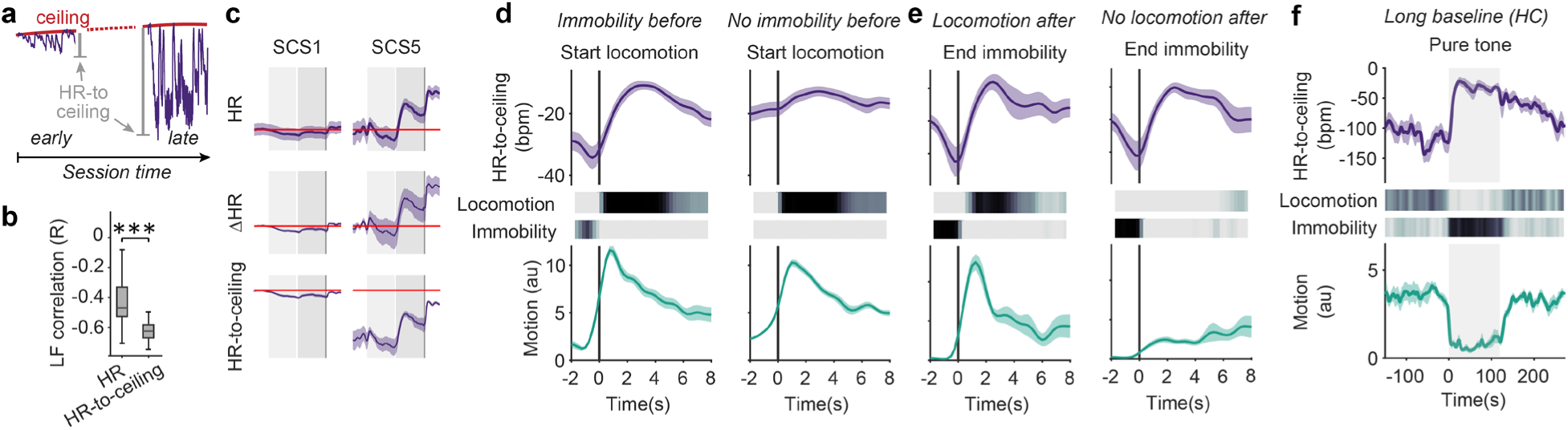
Defense state dynamics determine cardiac responses. **a,** HR-to-ceiling is the delta between the HR value at time *t* and the ceiling. Because of decreasing *rigidity*, the values appear greater further from the beginning of the recording. **b,** LF amplitude was only poorly correlated to raw HR, but well correlated with HR-to-ceiling (n = 33 mice; two-tailed Mann–Whitney U test). **c,** comparison of raw HR (top), ΔHR (middle) and HR-to-ceiling (bottom) average responses (n = 33 mice) for SCS1 (left) and SCS5 (right). The red line shows the baseline values before SCS1 to help comparison. While the dynamics are globally similar, raw HR and ΔHR suffer from higher variability, and encourage improper interpretations because their values do not capture an absolute level. **d,** PSTHs built around locomotion bouts start (n = 23 mice, OF), split in two groups depending on whether immobility was present shortly before the locomotion bout (left panel) or not (right panel; see also heat maps in the middle). In both cases, there is an increase in motion consistent with locomotion (bottom), and the HR-to-ceiling value during locomotion is similar (top). However, there is a clear increase in HR-to-ceiling only when transitioning from immobility to locomotion, not otherwise. **e,** PSTHs built around immobility bouts end (n = 23 mice, OF), split in two groups depending on whether immobility was directly followed by locomotion (left panel) or not (right panel; see also heat maps in the middle). HR-to-ceiling values increased in both cases, showing that the relevant state change is exiting immobility, and not engaging into locomotion. **f**, CS retrieval after a long baseline in the homecage led to a crisp immobility (probability heat map and motion curve, bottom; n = 8 mice), which was accompanied by a pronounced increase in HR (HR-to-ceiling, top). This highlights the dependency of the cardiac response not only on the stimulus and behaviour, but also on the conditions and context at a specific time point.

Additional analyses revealed that the marked differences between the two quantifications were due to a strong inter-dependency of microstates. When mice were immobile right before tone exposure, no further HR decrease was observed (Extended Data Fig. 2g,h), which further supported the idea that both evoked and spontaneous immobility/bradycardia microstates are a single entity, highlighted here by the continuum between the two. On the contrary, while locomotion bouts were associated with HR increases only when they were following an immobility bout (Fig. 2d), there was always an increase in HR at the end of an immobility bout, regardless of whether it was terminated by locomotion or non-locomotor behaviour (Fig. 2e). Similarly, the HR decrease following rearing could be attributed mainly to immobility bouts (Extended Data Fig. 2i). This not only shows that HR increases and decreases are not merely reflecting beginning and end of locomotion, respectively, but that entering or exiting the immobility/bradycardia microstate constitutes the decisive defensive state “switch”.

Importantly, our data also suggested that a HR decrease (or increase) *per se* is not specific for a certain microstate, but that the absolute distance to maximal HR at any given time is a major determinant of a microstate, best captured by HR-to-ceiling measure. To further probe the interdependency of micro- and macrostates, we conditioned another cohort of mice, and presented them with the CS (pure tone only) in their homecage, after a long baseline period. As expected, the long baseline allowed HR to return to low values and, remarkably, the CS presentation led to immobility as a behavioural response accompanied by a strong increase of HR (Fig. 2f, Extended Data Fig. 2j), paralleling earlier results using similar conditions^21, 30^.

These results show that cardiac responses, even for similar stimuli and during identical behaviours, depend heavily on the pre-existing state of the animal. Conversely, this demonstrates that cardiac changes differentiate between defensive reactions seemingly identical on the behavioural level.

### Context-dependency of cardio-behavioural defensive states

Consequently, we investigated whether cardiac states differentiate distinct microstates associated with stereotypical defensive behaviours in general. In fact, various defensive behaviours were related to significantly different levels of HR-to-ceiling values (Fig. 3a). Risk assessment behaviours such as stretch-attend posture and rearing were associated with the highest HR-to-ceiling values. Intermediate levels were found for locomotor behaviours, also confirming that increases in HR are not simply adjustments to physical needs^31^ (Fig. 3a). While raw HR allowed differentiation only between immobility and other behaviours, LF amplitude values identified more subtle differences albeit with less precision than HR-to-ceiling (Extended Data Fig. 3a,b). Because of the strong influence of *rigidity* changes over time on the HR responses and the intricate relation between behaviours and HR, we checked if one behaviour was predominantly expressed during, e.g. the initial period of high rigidity while another one would be rather present at the end of the recording, during low rigidity. However, temporal distribution of behaviours did not explain the differences observed (Extended Data Fig. 3c) and analyses of behaviour over time, while showing the effect of within-session *rigidity* development, confirmed the differences in average HR-to-ceiling values for the different behaviours (Extended Data Fig. 3d,e). As *ridigity* decreased, allowing for a wider range of HR values, differences between behaviourally defined microstates increased.

**Fig. 3.**
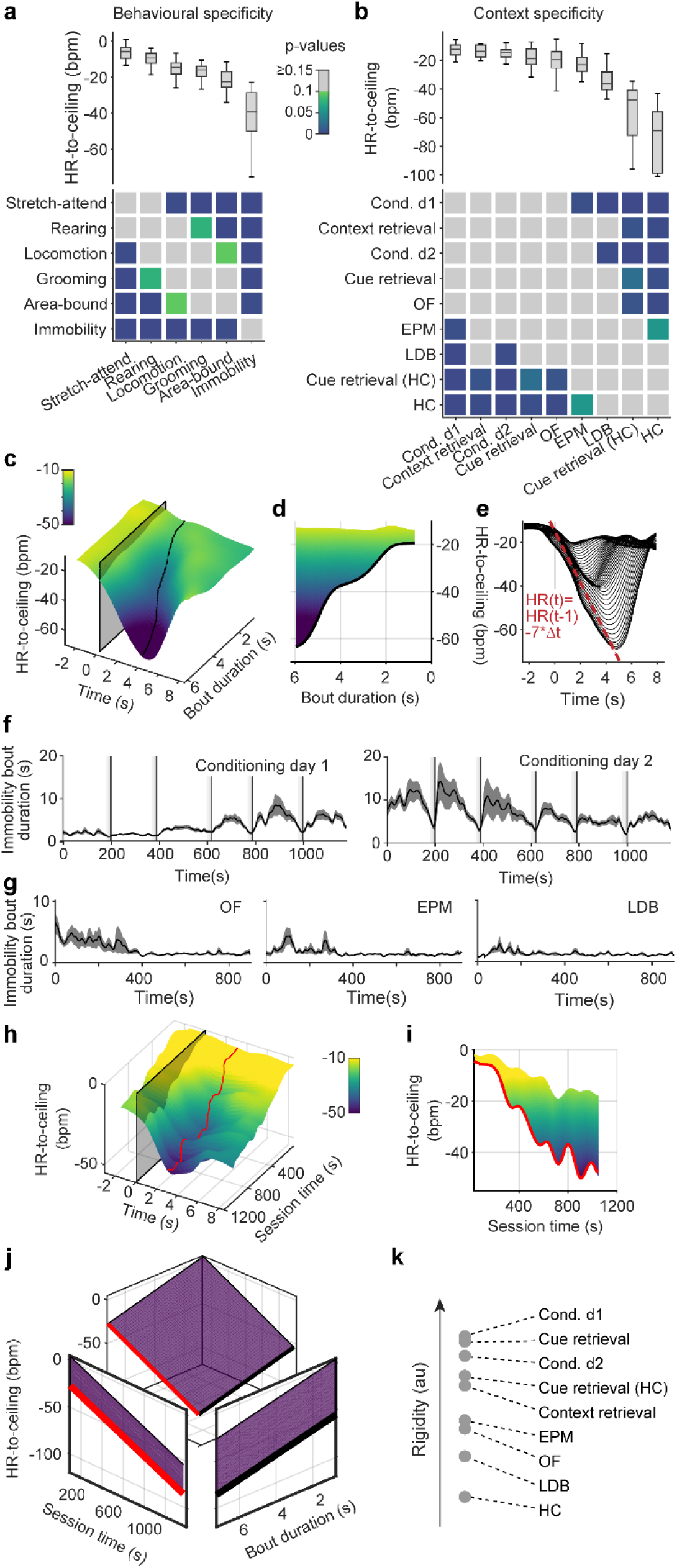
Individual defensive states depend on context and duration of behaviour. **a**, Average values of HR-to-ceiling for different behaviours during conditioning day 2 recordings (top), and grid showing the results of statistical evaluation of the differences between them (bottom; Kruskal–Wallis followed by post-hoc pairwise comparison with Bonferroni correction), showing a defined hierarchy (n = 33 mice). **b**, Comparison of the average values of HR-to-ceiling during locomotion in the different paradigms (top), and corresponding statistical analysis (bottom; OF: n= 23 mice; EPM: n = 20 mice; conditioning day 1: n= 30 mice; conditioning day 2: n= 33 mice; LDB: n = 12 mice; context retrieval: n = 5 mice; cue retrieval: n = 10 mice; cue retrieval (HC): n = 9 mice, HC: n = 10 mice; one-way ANOVA followed by post-hoc pairwise comparison with Bonferroni correction). High threat contexts are associated with higher values. **c**, 3D representation of the immobility-associated decrease in HR-to-ceiling during conditioning day 2 in function of bout duration (n = 33 mice), showing that longer episodes are associated with greater bradycardia. **d**, side view from **c** showing the correlation between immobility bout duration and bradycardia amplitude. **e,** 2D visualisation of **c**, showing the conserved kinetics across all bout durations, in particular for the decreasing portion (each line represents a range of bout durations values). **f-g,** Average immobility bout duration in function of time showing stereotypical changes for the different contexts (OF: n = 26 mice; EPM: n = 22 mice; LDB: n = 12 mice; conditioning day 1: n = 40 mice; conditioning day 2: n = 40 mice). **h-i,** 3D representation of the immobility-associated HR decrease during conditioning day 2 in function of the session time (**h**) and 2D side view (**i**), highlighting the dependency on the time within the session. **j,** fit integrating the amplitude of immobility-associated bradycardia in function of bout duration and time within the session (R² = 0.991). This allows to disentangle the contribution of the different factors, and to extract the influence of the session time, which would mainly reflect *rigidity*. Therefore, the curve in black is the equivalent of **d**, after accounting for the time within the session, while the curve in red is the equivalent of **i** after taking into account the influence of bout duration and its changing values in different contexts. **k,** values of the session time coefficients obtained from a similar analysis for the different paradigms. The coefficients ranking converges with the sorting of the contexts in terms of immobility- or locomotion-associated HR-to-ceiling values: the smaller HR deviations from the ceiling are correlated with a smaller relaxation over the session, overall reflecting a higher *rigidity*.

Next, we asked whether microstates reflect the behavioural context in which they occur. We therefore exposed mice to different threat contexts and focused on HR-to-ceiling values associated with locomotion, a behaviour present in all context conditions (Fig. 3b). Strikingly, ranked HR-to-ceiling values reflected expected contextual threat levels, with highest values under conditions of concrete threat (conditioned flight paradigm), intermediate values in paradigms with diffuse threat (Open Field, OF; Elevated Plus Maze, EPM; Light-Dark Box, LDB) and lowest values in the homecage (Fig. 3b). Again, raw HR yielded no differences while LF amplitude measures recapitulated the HR-to-ceiling mean values, albeit with much less variability for shock-related contexts (Extended Data Fig. 3f,g).

In contrast, ranking of contextual influence on immobility was less clear (Extended Data Fig. 3h). We reasoned that this was due to immobility bout kinetics and in fact found a strong correlation between the bradycardia amplitude and immobility bout duration during conditioning (Fig. 3c,d), with markedly stereotypical kinetics (Fig. 3e). In addition, we found that immobility bout duration differentially developed over time in different contexts: it progressively increased during initial threat conditioning day 1 (Fig. 3f, **left panel**), remained high on the second day of conditioning, but showed a decrease in the course of the session (Fig. 3f, **right panel**). Under diffuse threat conditions (OF, EPM, LDB), immobility bout durations were slightly longer at the beginning of the recording (Fig. 3g). This suggested that the interaction between both findings (dependency on bout duration, and stereotypical bout durations) could influence the results of indiscriminate averaging. In a comparable manner, the progressive changes in *rigidity* over time (Fig. 3h,i) certainly influence such quantification. Thus, taking into account the time- and context-dependency of immobility bout duration, we restricted them to certain ranges (last intertrial interval, ≤2.5s bout duration) to mitigate their influence. The result allowed ranking of the different contexts that resembled the context distribution for locomotion, but with comparatively reduced variability (Extended Data Fig. 3i).

This prompted us to find a way of integrating within the same analysis the two major influences, bout duration and time within the recording, on immobility-associated bradycardia. To this end, we extracted the peak bradycardic values for immobility episodes spanning the possible ranges of bout durations and time within the recording and fit a simple model to them (Fig. 3j, Extended Data Fig. 3k). The result was a linear equation with one coefficient (*A_t_*), accounting for within session changes in HR amplitudes, thereby accounting for the slow *rigidity* macrostate, and another for bout duration (*B_d_*). It followed that *Bradycardia_IB_* = *A_t_* ∗ t + *B_d_* ∗ *Duration_IB_*, with *Bradycardia_IB_* and *Duration_IB_*, i.e. the peak bradycardia and the bout duration of a given immobility episode, respectively, and *t* the time at which it occurred (Extended Data Fig. 3l; see also detailed explanations in Method section). By comparing the coefficients between the different paradigms, we were able to find a similar ranking as for the averages (e.g. Fig. 3b) for the session coefficients, which can be seen as a relative *rigidity* measurement (i.e., smaller coefficients mean that the time within the session is associated with moderate increase in bradycardia amplitude, and conversely). As such, paradigms with expected high threat levels (e.g. Conditioned Flight) had the lowest values, and therefore the highest *rigidity* overall (Fig. 3k).

We were therefore able to confirm that specific behaviours are associated with particular cardiac profiles, but also that the immobility-related microstate commonly termed ‘freezing’ is not a homogenous and static event determined only by its probability of occurrence. It crucially exhibits intrinsic behavioural properties, i.e. individual bout duration associated with dynamic cardiac changes. This is supported by the fact that immobility bout duration was not random, but instead exhibited context- and time-dependent patterns. Moreover, we show that expression of microstates is heavily influenced by slower underlying macrostates changes, which partially depend on the context in which they occur. Overall, this emphasizes their value in characterizing integrated defensive states.

### Integrated defensive states encode contextual threat levels

Our data demonstrated that HR exhibits high sensitivity to different threat levels induced by cues and contexts. Standard paradigms for fear and anxiety, solely developed based on behavioural readouts, use specific context designs to infer spatially-dependent threat levels^32^. We therefore asked next whether detrended HR-to-ceiling values would also report contextual subregions in classical tests evoking diffuse threats. Strikingly, we found a gradient of HR-to-ceiling values from low to high threat areas in elevated plus maze and light-dark box assays (Fig. 4a,b). Equivalent analyses based on raw HR did not yield any significant differences (Extended Data Fig. 4a,b). Surprisingly, no differences were observed between center and periphery of the open field test (Fig. 4c, Extended Data Fig. 4c), even though thigmotaxis and immobility were in line with standard expectations (Extended Data Fig. 4d-f). This unexpected result suggested that cardiac activity could reflect relative homogeneity (e.g. OF) or explicit compartmentalization (e.g. EPM, LDB) of threat contexts and prompted us to ask whether HR differences between certain subregions predicted the existence of state switches during transitions from one to the other context compartment. On the behavioural level, no such switch was visible, since mice exhibited locomotion patterns that differed in magnitude but not in direction when transitioning from low to high threat compartments and vice versa (Fig. 4d-f, **lower panels,** and Extended Data Fig. 4g-l, **lower panels**). In contrast, transitions from high to low threat areas were accompanied by marked decreases in HR, whereas transitions from low to high threat areas were concomitant with increases in HR, reflecting state switches mediated by the different subareas (Fig. 4d-f, upper panels, and Extended Data Fig. 4g-l, **upper panels**).

**Fig. 4.**
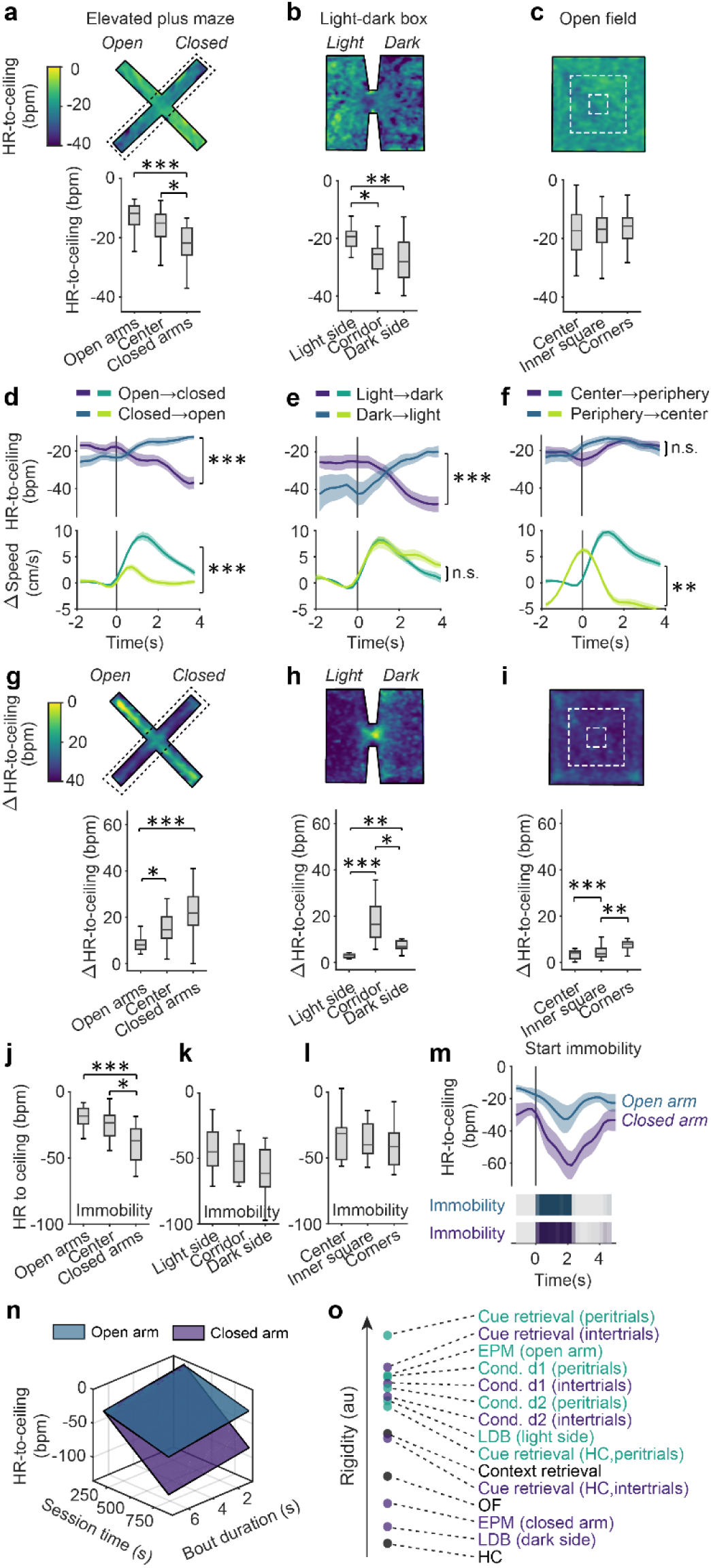
Integrated defensive states encode contextual threat levels. **a-c**, heat map representations of the mean HR-to-ceiling values in the different contexts (top) and corresponding quantification for discrete subareas (bottom; **a**, EPM: n = 20 mice, one-way ANOVA followed by post-hoc pairwise comparison with Bonferroni correction; **b**, LDB: n = 12 mice, one-way ANOVA followed by post-hoc pairwise comparison with Bonferroni correction; **c**, OF: n = 23 mice, one-way ANOVA). Average values are greater in high threat areas. **d-f**, PSTHs for transitions between low-threat and higher-threat subareas showing that the corresponding locomotion bouts (bottom curves) are associated with direction-specific HR-to-ceiling changes (top curves) in the EPM (**d**; n = 20 mice, two-tailed t-test) and LDB (**e**; n = 12 mice, two-tailed t-test), but not the OF (**f**; n = 23 mice, two-tailed Mann-Whitney test). t0 corresponds to the start of the area transition locomotion bouts. **g-i**, For each panel, heat map showing the average delta between highest and lowest HR-to-ceiling values expressed over the course of a recording in each position of the whole context (top), and the corresponding quantifications (bottom), highlighting a *rigidity*-like effect in certain areas, with reduced ranges (EPM: n = 20 mice, Kruskal-Wallis followed by post-hoc pairwise comparison with Bonferroni correction; LDB: n = 12 mice, Kruskal-Wallis followed by post-hoc pairwise comparison with Bonferroni correction; OF: n = 23 mice, Kruskal-Wallis followed by post-hoc pairwise comparison with Bonferroni correction). **j-l**, subarea-specific values of HR-to-ceiling during immobility (**j**, EPM: n = 20 mice, one-way ANOVA followed by post-hoc pairwise comparison with Bonferroni correction; **k**, LDB: n = 12 mice, one-way ANOVA; **l**, OF: n = 23 mice, one-way ANOVA). Immobility-related decrease in HR are of lower amplitude in high threat areas. **m,** PSTH illustrating that for immobility bouts of matching durations (bottom), the amplitude of immobility-associated bradycardia (top) is lower in the open than in the closed arms of the EPM. **n**, sheet analysis of the immobility-associated bradycardia integrating both bout duration and time within the session for closed *vs* open arms (respectively R² = 0.989 and R² = 0.97), confirming a globally smaller amplitude in the open arms (p<0.001, n = 20 mice). **o,** Values of the session time coefficients from sheet analyses performed on the different contexts, differentiating between high *vs* low threat subareas or periods. Higher threat is associated with smaller coefficient, uncovering a threat level-related macrostate characterized by higher relative *rigidity*.

What is the nature of these context-dependent states? We hypothesized that contextual threat levels evoked a macrostate mechanistically similar to *rigidity*, i.e. imposing a restriction of HR-to-ceiling values in high-threat areas. This became apparent when instead of taking mean values per bin, deltas between maximum and minimum of the detrended HR-to-ceiling (Fig. 4g-i, Extended Data Fig. 4m-r) were used to identify subarea-specific changes. Decisively, smallest HR ranges are apparent in high-threat areas, such as open arms in the EPM, light side in the LDB, and centre in the OF (Fig. 4g-i). These findings were reminiscent of the contracted HR variability, i.e. the *high rigidity* macrostate at the beginning of a threat conditioning session (Fig. 1d-g). We therefore looked at the amplitude of immobility-associated bradycardia across the different subareas, which was significantly smaller in the high threat areas (EPM open arms, LDB light side) than in low threat areas (EPM closed arms, LDB dark side; Fig. 4j-l) mimicking changes observed during fear conditioning. Crucially, this did not rely on different intrinsic behavioural properties, as it stood true even when we matched immobility bout duration (Fig. 4m, Extended Data Fig. 4s). Applying the integrated analysis of immobility-associated bradycardia confirmed this finding, as open and closed arms presented very distinct profiles, with higher rigidity observed in the open arm (Fig. 4n, Extended Data Fig. 4t). We hypothesized that this could be a general property of HR that would reflect threat level, and performed a similar analysis on the different paradigms after differentiating low *vs* high threat periods (i.e. inter-CS intervals *vs* peri-CS intervals), or low *vs* high threat areas as for the EPM. The updated ranking met those expectations, demonstrating that rigidity globally reflects threat level that in turn affects its development over time (Fig. 4o, Extended Data Fig. 4t). Together, the integrated defensive state analysis reflected subarea-specific differences of threat level in classical fear and anxiety tests.

### *Chx10*-positive neurons in the midbrain periaqueductal grey mediate a cardio-behavioural defensive microstate

If such ubiquitous interactions between micro- and macrostates define defensive states on the level of the associated responses, we reasoned that they could help to pinpoint neuronal circuit elements mediating defensive states, such that the evoked responses should present similar properties as in the naturally occurring reactions. We therefore used optogenetic manipulations of specific neural circuit elements involved in mediating defensive behaviours. Previous studies assigned such a role to glutamatergic neurons in subregions of the midbrain periaqueductal grey (PAG), a key region for the defense reaction, and supported a labelled line-like mechanism of functionally defined PAG output pathways^33, 34^. Glutamatergic neurons, characterized by the expression of vesicular glutamate transporter 2 (*Vglut2)* of the lateral and ventrolateral PAG mediated defensive behaviours, ranging from flight to threat-induced immobility, i.e. freezing, the latter associated specifically with glutamatergic ventrolateral PAG neurons projecting to the medullary magnocellular nucleus (Mc) and expressing *Chx10*^33–35^. Importantly, these studies left unanswered the question of whether and how these circuit elements take part in mediating the integrated defensive response, a role that has long been postulated for PAG circuits^6–8, 36, 37^.

We equipped naive *Vglut2-Cre, Chx10-Cre*, and *Vglut2-Flp/Chx10-Cre* mice with ECG electrodes (Fig. 5a), and locally injected an adeno-associated viral vector (AAV) into the vlPAG to Cre-dependently express ReaChR (Fig. 5b and Extended Data Fig. 5a), a red-light activatable, excitatory optical actuator^38^. Stimulating *Vglut2+* neurons evoked intensity-dependent responses: low intensities led to immobility and bradycardia, while higher intensities led to mixed flight and transient immobility as well as bradycardic HR responses (Fig. 5c). Optical activation of the *Chx10+* population resulted in robust immobility and bradycardia (Fig. 5d). This strongly suggested that *Chx10+* neurons are a subgroup of glutamatergic vlPAG neurons mediating a particular defensive microstate, i.e. immobility concomitant with bradycardia. Confirming this distinct role, specific optical activation of *Vglut2+/Chx10-* via a double-conditional, intersectional approach recapitulated the behavioural activation seen when activating the entire *Vglut2+* neuronal population, but failed to produce any cardiac effect (Fig. 5e). This was in contrast to HR decreases evoked by optoactivation of the entire Vglut2+ neuron population or the tachycardic response associated with spontaneous flight (Extended Data Fig. 5b). While activation of all glutamatergic vlPAG neurons with high intensity resulted in a rather non-natural state of strong locomotor behaviour concomitant with HR decrease, the *Chx10*-mediated effect resembled the immobility-bradycardia microstate we had characterized before (**see** fig. 1f). Using different optogenetic protocols first allowed us to confirm that the amplitude of the evoked-bradycardia did in fact increase with stimulation duration, with the same relationship as in spontaneous episodes derived from animals which underwent an OF test without optogenetic perturbation (Fig. 5f and Extended Data Fig. 5d). Inhibition of *Chx10+* neurons during a context threat memory retrieval blocked sustained immobility (Fig. 5g and Extended Data Fig. 5e). To add further construct validity to these findings, we asked whether the evoked and spontaneous microstate were similarly affected by *rigidity*. Across repeated optical stimulation of same length and intensity, the amplitude of the evoked-bradycardia linearly increased in function of time from the first to the fifth trial (Fig. 5h and Extended Data Fig. 5d), demonstrating that like the natural immobility bradycardia microstate, the evoked microstate was sensitive to *rigidity*. These data obtained by targeted perturbation of PAG circuit elements within our defensive state framework characterize *Chx10+* neurons as generators of an integrated defensive microstate, characterized by immobility and bradycardia.

**Fig. 5.**
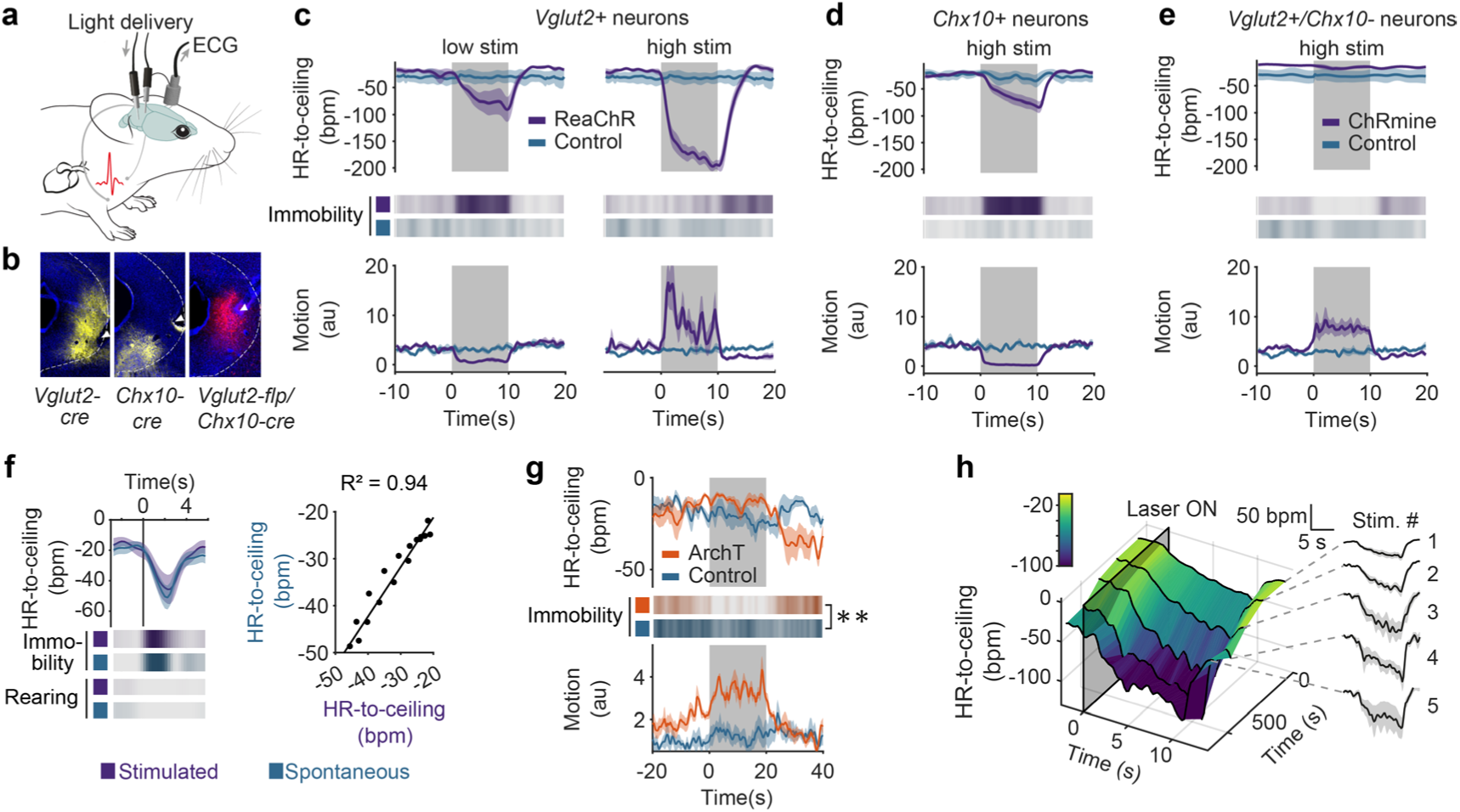
Specific circuit elements in the midbrain periaqueductal grey mediate integrated defensive microstates. **a**, ECG are recorded in freely moving mice while light stimulations are delivered to optogenetically activate specific vlPAG neuronal populations. **b,** Representative microscope images showing opsin expression and optic fiber placement for vlPAG optogenetics experiments. **c,** Optogenetic stimulation of *Vglut2+* neurons in the vlPAG led to different microstates depending on the stimulation intensity. Both caused a pronounced decrease in HR (top curves), but while the low intensity stimulation produced immobility (probability plot, middle), also visible via a decrease in motion (bottom curve), the high intensity stimulation led to the expression of active behaviours instead (both n = 8 vs controls, n = 5; two-tailed t-test for low intensity, Mann-Whitney U test for high intensity). **d,** Light activation of vlPAG *Chx10+* neurons evoked a defensive reaction resembling the one evoked by low-intensity stimulation of glutamatergic neurons (**c**) and spontaneous episodes, characterized by bradycardia and immobility (n = 8 vs controls, n = 5). **e,** ChRmine-mediated activation of the *vGlut2+/Chx10-* neurons in the vlPAG recapitulated the behavioural activation seen during high intensity stimulation of the global *Vglut2+* population, but without any bradycardia. **f,** Bradycardic responses evoked by optogenetic stimulation of *Chx10+* cells (n = 8 mice) present the same dynamic and amplitude as spontaneous immobility episodes (left curves) with similar immobility bout duration (heat maps, middle), as shown by the correlation (right, correlation of the time-matched HR-to-ceiling values during immobility between spontaneous and evoked episodes). **g,** Optogenetic inhibition of the *Chx10+* neurons interrupts context retrieval-driven immobility, suggesting again their involvement in the naturally occurring defensive immobility bouts (n = 5 vs controls, n = 4; two-tailed t-test). **h,** 3D representation of the amplitude of HR decrease evoked by optogenetic stimulation of the *Chx10+* neurons (n = 8 mice) in function of the relative time from the beginning of the recording (left), showing a progressive increase, also visible on the single PSTH curves (right), and suggesting it is sensitive to the *rigidity* macrostate in a similar manner as spontaneous immobility episodes.

### Multidimensional analysis reveals overarching defensive states

Using behavioural and cardiac autonomic readouts enabled us to define integrated defensive micro- and macrostates. To ensure generalisation and translation of this cardio-behavioural framework, we next aimed at integrating multiple parameters derived from behavioural and cardiac readouts commonly used in fear and anxiety research. We therefore selected four variables, reflecting the main states present in our conditions (i.e. HR-to-ceiling, immobility bout duration, immobility-associated bradycardia, LF amplitude during locomotion), for which we processed average values in function of time for each paradigm. Subsequent ‘Uniform Manifold Approximation and Projection for Dimension Reduction’ (UMAP^39^) showed that different behavioural paradigms cover different 2D (Fig. 6a) and 3D (Extended Data Fig. 6b) subspaces, a phenomenon also visible between the open and closed arms of the EPM (Fig. 6a,b and Extended Data Fig. 6a-d). These findings demonstrate that combined analysis of multiple readouts allow capture of a general defensive state, associated with a specific context, time, or both.

**Fig. 6.**
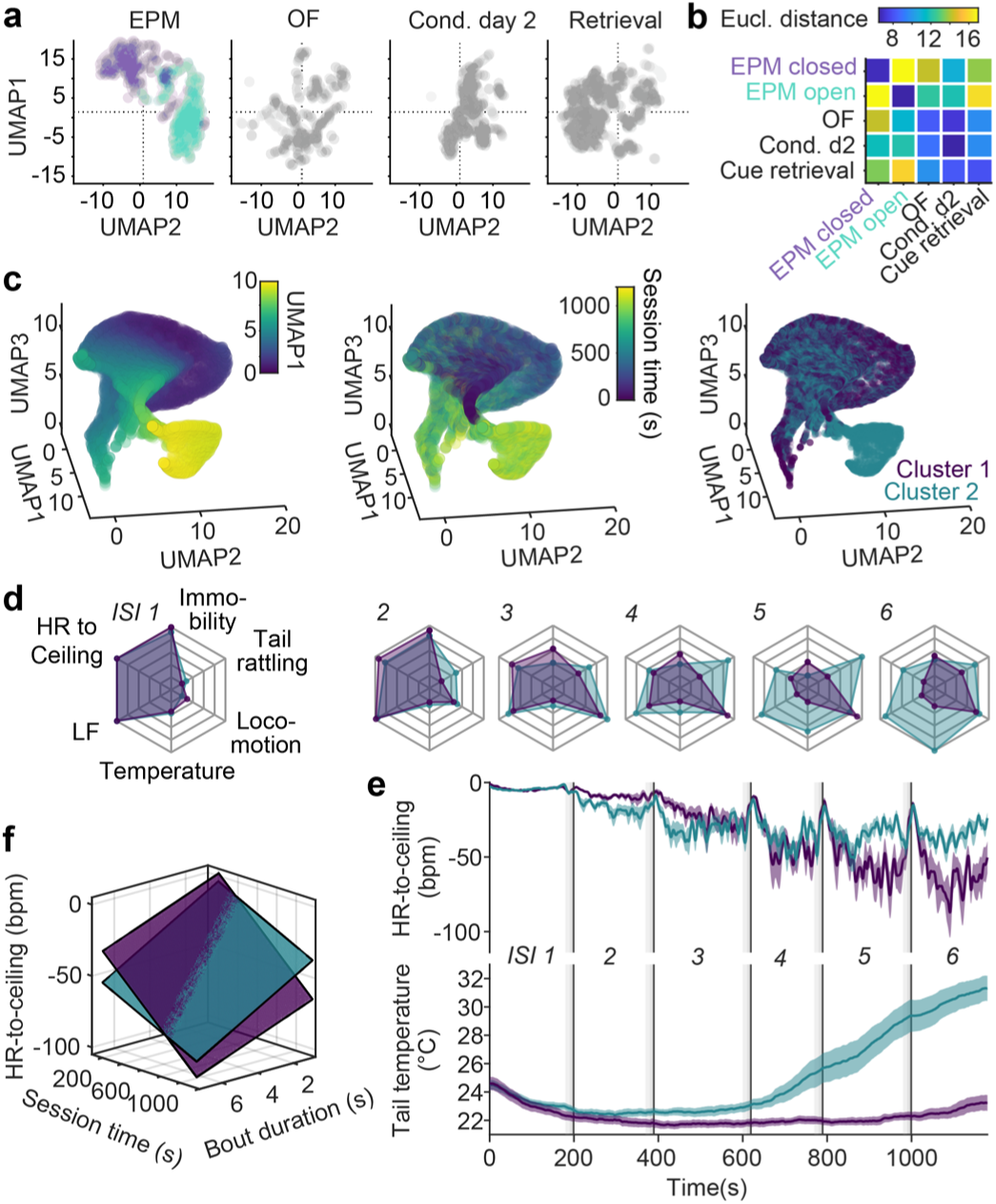
Multidimensional analysis reveals overarching defensive states. **a**, 2D representation of the UMAP dimensionality reduction of secondary readouts of interest HR-to-ceiling, immobility bout duration, immobility-associated bradycardia, LF amplitude during locomotion) extracted for different paradigms. The two-coloured sub-clouds for the EPM correspond to the open arm and closed arm data points. **b**, Matrix of the average Euclidean distances between the different contexts corresponding to the UMAP results in **a**, informing about states space relative proximity (average of 250 UMAP runs). **c**, Exploratory 3D plots of UMAP dimensionality reduction applied to a set of raw readouts from conditioning day 2 (including tail temperature) suggested the existence of a subcluster (left, colour coding for UMAP dimension 1 for better readability), which seemed to coexist with a main cluster later during the recordings (middle). The results from a BIC-guided k-means clustering of mice averages produced two clusters of mice that corresponded to the previously suspected clusters (right). **d**, Spider charts showing the average values of several readouts over the 6 ISI for the two groups of mice identified (respectively n = 14 vs n = 15), with progressively pronounced autonomic and behavioural differences. **e,** Average curves of HR-to-ceiling and tail temperature, corresponding to the clusters visible in **c**. The higher average values of HR-to-ceiling from the mid-recording on in the group with an increase in tail temperature suggests a higher *rigidity*. **f,** Sheet analysis confirms that the cluster of mice with an increase in tail temperature presents an overall higher rigidity, with different dynamics (cluster 1, no temperature increase, R² = 0.983, n = 14; cluster 2, temperature increase, R² = 0.98, n = 15; comparison, p<0.001). This underlines that added readouts can help identifying further underlying integrated macrostates.

Because defensive states encompass adaptive changes beyond cardio-behavioural dimensions, we lastly tested whether additional state dynamics could be revealed by integration of more autonomic and raw-data readouts, without making use of our understanding of some of the underlying states as before. Therefore, we performed UMAP analysis of speed, motion, HR-to-ceiling, LF amplitude, as well as thermographically recorded tail temperature obtained during the conditioned flight paradigm. Interestingly, a sub-cluster was visible on the initial view (Fig. 6c, **left**), which seemed to develop over time of the behavioural session (Fig. 6c, **middle**). We used the Bayesian Inference Criterion (BIC) to determine whether averages per mouse could differentiate groups, followed by k-means clustering. This procedure identified two clusters (Fig. 6c, **right**) corresponding to two groups of mice that diverged particularly regarding their thermoregulatory and cardiac profiles (Fig. 6d,e and Extended Data Fig. 6e-g). Integrated analysis of the immobility-associated bradycardia confirmed major differences in terms of *ridigity* development over the course of the session, with the group of mice displaying an increase in tail temperature presenting limited decrease of the *rigidity* over time, thereby pointing towards putative macrostate dynamics dependent on inter-individual differences (Fig. 6f). Overall, multidimensional analysis extended our previous findings (**see** Fig. 4) by revealing overarching contextual macrostates associated with different behavioural paradigms and characterized by inter-individual differences in state dynamics.

## Discussion

Our data demonstrates that defensive states, as previously and intuitively postulated, consist of integrated and interdependent processes. Based on the detailed characterization of the technically simple readout HR and state-of-the-art behavioural analysis, we developed a framework that integrates behavioural and cardiac components and reconciles the seemingly contradictory findings on threat-induced HR changes. We here gather strong evidence of the existence and interaction of short-lasting *microstates* and *macrostates* with slower dynamics as important determinants of defensive states.

Our work reconciles the long-standing dispute in the field of whether threat exposure in general, and the immobility state termed freezing in particular, is associated with tachy- or bradycardia. We show that both responses occur and that they crucially depend on context-related internal macrostates such as *ceiling* and *rigidity* as well as the preceding microstate. The initial responses to threat are macrostate changes that promote overall tachycardia. On the background of this long-term change, an important defensive microstate is characterized by immobility and bradycardia. While this microstate is induced by contextual as well as cue-induced threats, composition of its behavioural and cardiac components greatly differs with respect to immobility bout duration and HR-to-ceiling values. This strikingly reflects the influence of macrostates on a readout most commonly used as a single entity to quantify fear, i.e. ‘freezing’ behaviour^40–42^. As a result, our data demonstrates the existence of different defensive states associated with similar readouts such as immobility or HR decrease. When binarizing defensive states into freezing vs. non-freezing periods, as well as when integrating over long periods of time (e.g. by using average freezing values), insights on the important correlations of micro- and macrostates get lost, thereby preventing potential attributions of neuronal activity with a particular readout. On the other hand, taking into account the integrated and dynamic nature of the threat-related immobility, commonly termed as the freezing defensive state, allows for clear attribution of the underlying circuit elements, which are excitatory *Chx10+* neurons within the vlPAG. This shows how combining classic gain- and loss-of-function approaches with previously identified micro- and macrostates properties can strengthen the interpretation of a certain circuit element being involved in a natural state. Overall, our findings refine the ‘labelled line’ concept^4, 9, 43, 44^, which postulates the existence of functionally defined neuronal pathways, by addressing the multitude of highly dynamic and interacting brain functions in the context of defense against threats.

Our results reveal that for several integrated defensive states, the average distance to an approximated maximal HR (HR-to-ceiling) is the most relevant characteristic. This suggests that physiologically, there is not a “fixed” signal systematically leading to a HR change of similar amplitude, but that HR is adjusted to a particular set point for individual microstates associated with specific defensive behaviours. Importantly, these microstate-related set points are affected by the (sub)context-dependent changes in the *rigidity* macrostate. Our framework includes discretizing HR data to unravel semi-stable states, but also takes into account temporal dynamics, which can characterize states of their own. This is most evident in the described *ceiling* and *rigidity* macrostates, capturing the upper boundary and range of cardiac output, respectively. Importantly, while *ceiling* is predominantly determined by cardiac function and thus may reflect sluggish peripheral endocrine processes, the richer dynamics of *rigidity* report interaction of cardiac and behavioural defensive states. Our finding that LF oscillations, which have been associated with emotions in humans^45^ are sensitive to *rigidity* strongly suggests that these two processes share an underlying mechanism, such as baroreceptor reflex function. *Chx10* cells of the vlPAG, while serving as microstate generators, do not affect *rigidity* and thus are unlikely to regulate baroreflex function directly. Thus, it will be important in the future to determine how brainstem and higher-order circuits control temporal dynamics of central and peripheral macrostates. Nervous systems have been modeled as near-chaotic systems stabilized by attractors^46^. Our data suggests similar state dynamics, i.e. main and partial attractors as macro- and microstates, respectively, for behavioural and cardiac readouts. It is conceivable that this is closely related to fractal characteristics of HR, which exhibit rigidity after pharmacological stimulation^47^. Overall, it reaffirms our framework’s adequacy in providing a window into higher-order brain functions such as defense against threat. Importantly, defensive state dynamics involve bi-directional brain-body communication, i.e. somato- and visceromotor control as well as interoception^48^.

From a methodological perspective, we show that when solely relying on behavioural readouts, the complexity of “hidden” state dynamics is lost. Moreover, HR can indicate distinct threat-dependent state shifts even when behaviour is redundant. However, higher-order defensive states derived from human emotions such as ‘fear’ and ‘anxiety’ have classically been quantified in behavioural paradigms validated by pharmacology^49^. Our framework, beyond identifying defensive states in standard behavioural assays used to model fear and anxiety, demonstrates that context is a major determinant of defensive microstates, which appear identical on the behavioural level, but exhibit different cardiac components. In addition, our data reveals large differences between various defensive macrostates in contexts that explicitly provide safe environments (EPM and LDB) and a uniform macrostate elicited in those that do not (OF).

We show that analyses of biologically relevant read-outs of different modalities allows differentiating behaviourally similar group differences. Ideally, unsupervised clustering algorithms should be used to identify subgroups within a global “state space”, as a basis for identifying novel biomarkers for normal as well as pathological state dynamics. In taking a step back with a stringent data-driven approach, we provide a framework for characterization of multimodal, integrated defensive states, thereby reopening a pathway towards translational research across species and from normal to maladapted fear and anxiety.

## Methods

### Animals

2-to 6-months old C57BL/6 male wild type as well as transgenic mice were bred in an in-house animal facility. Slc17a6tm2(cre)Lowl (Vglut2-ires-Cre) and Gad2^tm2(cre)Zjh^ (Gad2-ires-Cre) knock-in mice were initially obtained from Jackson Laboratory (respectively stock number 028863 and 010802). Chx10-ires-Cre mice were bred from parents kindly provided by Prof. Ole Kiehn (University of Copenhagen)^50^. Slc17a6tm1.1(flpo)Hze (Vglut2-ires2-FLPo-D) knock-in mice, obtained from Jackson Laboratory (stock number 030212) and Chx10-Cre mice were cross-bred in-house. All mice were individually housed on a 12h/12h light-dark cycle and experiments were performed during the light cycle. Food and water were available *ad libitum*. All animal procedures were approved by the local veterinary authorities and animal experimentation ethics committee (Regierung von Unterfranken, authorization 2532-2-509). Present reporting follows ARRIVE guidelines (Animal Research: Reporting of In Vivo Experiments)^51^. Sample sizes were estimated based on previous studies using similar experimental designs^24, 33^.

### Stereotactic surgeries

Adeno-associated viruses (AAVs) were used as recombinant viral vectors to deliver genetic constructs of interest. Viruses for optogenetic experiments were produced in an in-house viral production facility from plasmids ordered from Addgene (pAAV-hSyn-flex-ReaChR-Citrine #50955, pAAV-hSyn-DIO-mCherry #50459) or ordered ready-made from Addgene (pAAV-FLEX-ArchT-tdTomato #28305, pAAV-nEF-Coff/Fon-ChRmine-oScarlet #137160). Mice were anesthetized with isoflurane (Harvard Apparatus, Iso-Vet Chanelle) in O2-enriched air (induction: 4%, maintenance: 1.5-2%). For systemic intra-operative analgesia, buprenorphine (0.05-0.1 mg/kg; Buprenovet, Bayer) was administered subcutaneously 20 minutes prior to the start of the surgery. During the surgery, body temperature was kept stable by keeping the animal on a heating pad. Mice were fixed in a stereotactic frame (Model 1900, Kopf) and local analgesia was achieved by injecting ropivacaine under the scalp (8 mg / kg, Naropin, AstraZeneca), after which a midline incision was performed. For viral injections targeting the vlPAG, craniotomies (0.4 mm diameter) were drilled 4.6 mm caudally and ±0.5 mm laterally from bregma. A calibrated glass pipette (calibrated micropipette 1-5 µl, Drummond Scientific) filled with the appropriate virus was slowly lowered to the target depth of 2.9 mm below bregma. A volume of 70-100 nl was injected at a speed of 25 nl / min with a pressure injector (PDES-02DX, NPI electronic). After injection, the capillary was hold in place for 10 minutes, before being slowly withdrawn. The wound was sutured and treated with antiseptics. To ensure post-operative recovery analgesia, meloxicam (5 mg / kg every 12 - 24 hours; Metacam, Boehringer Ingelheim) was administered subcutaneously after the operation was completed. For optogenetic light delivery to the target brain region, custom-built optical fiber stubs (ceramic ferrules, Thorlabs; multimode fiber 0,39 NA 200 µm, Thorlabs) were implanted in a second surgery. Salp was opened by removing a skin patch of 0.7 x 0.7 mm and skull was cleaned and carefully dried. Craniotomies were performed 4.6 mm caudally and ±1.6 mm laterally to bregma. Fiber stubs were inserted in a 20° angle to a depth of -2.1 mm from brain surface and secured with cyanoacrylate glue.

### Measurement of the electrocardiogram (ECG)

To measure heart rate parameters, all animals were implanted with ECG electrodes. For this purpose, three PFA-coated wires (7SS-1T, Science Products) were soldered onto micro connectors (A79108, Omnetics, MSA components). Two wires were used to record the ECG signal differentially, the third was used as reference. During the surgery two small incisions were made on the upper right and lower left torso, at the ends of an imaginary diagonal centered on the heart. Wires were threaded through a blunt feeding needle in a subcutaneous tract carefully created from the skin incision on the front side of the mouse towards the scalp opening. The distal ends of the wires were stripped of their insulation over 3 mm and sutured onto muscle tissue of the thorax. The skin was closed, disinfected and the connector was fixed on the skull.

### Optogenetics

After a recovery period of at least one week after surgery, animals were handled four days prior to the first recording session in order to habituate them to the connection procedure. For optical stimulation, a LED fiber light source was used (Ce:YAG optical head, Doric; 582 nm bandpass filter for ArchT and ChRmine and 612 nm bandpass filter for ReaChR). Stimulation protocols were created in Radiant (Plexon), and an analog output from PlexBright (Plexon) was used to control stimulations’ intensity and patterns. For naïve testing of optogenetically-evoked effects, animals were placed into cylindric contexts (30 cm diameter), the floor being covered with homecage bedding. Vglut2-ires-Cre as well as Vglut2-ires2-FlpO/Chx10-Cre animals animals were stimulated 5 times for 10 s at either low (0.3-0.5 mW, constant) or high intensities (3.5 mW, 10 Hz, 20 Hz or constant) in 2 separate sessions. Chx10-cre animals, similarly received 5 times 10 s stimulations at 30 Hz stimulation frequency (7.2 mW). For short stimulations (1.1 and 2 s alternating, stimulation settings see above) of Chx10-Cre and long stimulations of Gad2-ires-Cre animals (5 times for 30 s, 12 mW, 30 Hz), subjects were placed into an Open Field context (50 x 50 cm) without bedding. For loss-of-function experiments, Chx10-Cre animals were conditioned with the Conditioned flight paradigm (see below). After two days of conditioning, animals were placed back to the conditioning context for 15 minutes and stimulated 4 times for 20 s (12 mW, constant).

### Histology and microscopy

Mice were anesthesized with a mixture of Ketamin (100 mg/kg) and Xylazine (10 mg/kg), injected intraperitoneally, and transcardially perfused with 1 x PBS and 4 % paraformaldehyde (PFA) for 5 minutes each. Brains were dissected and post-fixated over-night in PFA at 4°C. Samples were washed, embedded in 6% agarose cubes and cut into 60 µm coronal sections using a vibratome (Leica VT1200). Sections of Vglut2-ires2-FlpO/Chx10-Cre animals were immunhistochemically stained (primary antibody: rabbit anti-RFP, Rockland, 600-401-379; secondary antibody: donkey anti-rabbit Cy3, Jackson Immuno Research, 711-165-152). Sections were incubated in DAPI (4′,6-diamidino-2-phenylindole), mounted onto object slides and embedded with custom-made glycerol-based medium (Fluorostab) before imaging with a fluorescence microscope (AxioImager 2, Zeiss).

### Behaviour recordings

Behavioural experiments were conducted in two different sound-attenuated chambers (length: 100 cm, width: 80 cm, height: 116 cm) lit from above by an adjustable circular LED lamp (LED-240; Proxistar). One chamber was covered with white insulating foam, brightly lit (350 Lux), and contained a Petri dish filled with 70% ethanol (chamber E), while the other was covered with black insulating foam, dimly lit (130 Lux), and contained a Petri dish filled with 1% acetic acid (chamber A). For the different paradigms, the appropriate contexts were placed in the center of either of the chambers, and were cleaned with the corresponding liquid before each recording (either ethanol or acetic acid). Context temperature was kept at 22.5±1 °C. Different paradigms were used in order to capture as many different states as possible.

### Homecage (HC)

Animals were recorded in their homecage and left unperturbed for 15 minutes.

#### Open field (OF)

The apparatus consisted in a 50 cm x 50 cm x 50 cm white box. Animals were placed inside the middle of the box and left to freely explore the environment for 15 min.

#### Elevated plus maze (EPM)

The apparatus consisted of two open and two enclosed arms (8 cm width, 28 cm arm length, 28 cm wall height), elevated 25 cm above the chamber floor. Animals were placed in the crossing area at the intersection of the four arms, facing an open arm, and left to freely explore for 15 minutes.

#### Light-dark box (LDB)

The context box consisted of two compartments of identical dimensions (30 cm x 15 cm) that communicated via a 5 cm opening. One side was made of white material, with LED strips fixed near the upper rim providing illumination restricted to that compartment (350 LUX), while the other was black and void of any light source (50 LUX). Animals were habituated to dim light 15 minutes prior to the experiment.

#### Conditioned flight paradigm

The conditioned flight paradigm^24^ is a Pavlovian fear conditioning paradigm in which a serial compound stimulus (SCS) is used as the conditioning stimulus (CS) that is paired with the unconditioned stimulus, a shock. The SCS consists in a 10-s pure tone period (7,5 kHz, 75 dB, 500 ms beeps, 1 Hz) followed by a 10-s white noise period (1-20 kHz, 75 dB, 500 ms bursts, 1 Hz). On the first day (pre-exposure), the SCS alone is presented four times in a white cylinder (27 cm diameter). The conditioning was then performed on two consecutive days, with five SCS /US pairings on each day after a 3-min baseline period (pseudorandomized inter-stimulus interval (ITI): 170-230 s). The conditioning context was a red transparent square box (30 x 30 cm) with a grid floor through which footshocks were delivered. For each SCS/US pairing, a 1-s footshock (0.9 mA; Model 2100 Isolated Pulse Stimulator, A-M System) immediately followed the last white noise burst. On the 4th day, animals underwent cue retrieval in a transparent cylinder (30 cm diameter), to receive 16 SCS presentations, without US pairings (pseudo-randomized ITI: 80-140 s). One group of mice were placed into their homecage while replaying the SCS tones instead a new context. A subset of mice underwent an additional day of recording and were placed back in the conditioning context for 15 min on the 5th day of the protocol (context retrieval).

Days 1 and 4 of the conditioned flight paradigm (pre-exposure and cue retrieval) as well as OF, EPM, and LDB recordings were performed in context A (acetic acid smell, dim light), while recordings for days 2, 3 and optionally 5 of the conditioned flight paradigm (conditioning, context retrieval) were made in context E (ethanol smell, brighter light).

#### Retrieval in Homecage with long baseline

Animals were conditioned as described above. On the fourth day (retrieval), animals were placed in their homecage and left unperturbed for 40 minutes. The pure tone component (7,5 kHz, 75 dB, 500 ms beeps, 1 Hz) of the SCS was played for two minutes. The total recording time was 50 minutes.

### Recording system

The overall recording system consisted of several main elements. An acquisition system (Plexon, Omniplex system) recorded analog as well as digital signals and was coupled with CinePlex Studio (Plexon) for synchronized top RGB camera recordings (Pike Camera F-032C, Allied Vision, Campden Instruments). The Radiant software was used to create optogenetics stimulation protocols and control the global timing of experiments via PlexBright analog and digital outputs. An RZ6 multi-processor (Tucker-Davis Technologies) was used to deliver acoustic stimulations via a multi-field magnetic speaker (MF1, Tucker-Davis Technologies), control the shocks delivered by the stimulus isolator, and overall provide online processing and synchronization via a MATLAB/ActiveX/RPvdsEx interplay (MATLAB2019b, The MathWorks; RPvdsEx, Tucker-Davis Technologies). In particular, a fast initial train of TTLs followed by a 1Hz signal was generated and broadcasted to the different systems for offline alignment. Temperature data was acquired with a long wavelength infrared camera (A655sc, FLIR), via FLIR’s SDK within MATLAB. A custom GUI was used for pre-recording calibration and focus, and to visualize the movie in real-time. Recordings were triggered by MATLAB after the running signal was broadcasted by PlexBright and received by the RZ6, and similarly stopped. Thermal data was directly saved into a .seq file, and MATLAB was periodically saving the current number of acquired frames which allowed for offline synchronization. ECG data was acquired at 5 kHz via an amplifier (DPA-2FX, npi) connected to the OmniPlex system. Depending on the quality of the signal for the two electrodes, signal was saved differentially or from single electrodes.

### Heart rate analyses

Detection and extraction of heart beats from the raw ECGs was performed within a MATLAB GUI (custom code). Briefly, the raw signal was read from the .pl2 files using Plexon’s SDK. When needed, bandpass filtering was applied after adjusting the frequencies. The resulting signal was then raised to the 4th power to increase separation and smoothed with a Gaussian filter. A threshold was then defined to extract putative heart beats. After the timestamps were obtained from the modified signal, putative heart beat waveforms were extracted from the signal. A divergent template-matching and interbeat interval confidence scoring algorithm was used to pre-treat the results, with a high specificity. Uncertain ranges were left to be manually fixed by the experimenter. If that was impossible or there was any doubt because of a bad signal-to-noise ratio (e.g. contamination by electromyogram), the concerned ranges were marked, and excluded from further analyses. The R peaks were then extracted from each waveform, and saved for further processing.

#### Heart rate and wavelets

Heart rate was processed from the R peaks using a sliding window of 0.6 s ending at each peak, and resampled from these R peaks-based times to a fixed sampling rate. Continuous 1-D wavelet transform (MATLAB wavelets toolbox, The MathWorks) was used to extract the frequency band of interest (0.4-0.8 Hz), which corresponds to what has been hypothesized to be the murine equivalent of the LF band of HRV^45^ and the 0.1 Hz Mayer waves in humans^29^, closely related to baroreflex function.

#### Theoretical maximum heart rate (“ceiling”)

The local theoretical maximum HR curve was obtained from a resampled (4Hz) and median filtered HR (sliding window of 4 samples, centered around each sample) for the whole recording, by first extracting local maxima (sliding window, with more samples in the backward direction), and then smoothing with a sliding quadratic linear regression (‘smoothdata’ function in MATLAB, with ‘loess’ as method).

### Mouse tracking and behaviour scoring

Raw top view movies were processed with custom MATLAB code. For RGB movies, the 80th percentile of a manually or automatically selected set of frames was used as background that was substracted from all the frames. Then both for RGB and thermal movies, a threshold was manually selected on a GUI, and the resulting binary mask underwent a series of simple treatments: morphological closing, removal of small objects, another morphological closing, and finally morphological opening. The values for the different parameters were set to accommodate the different conditions. For each frame, mouse contour and center of gravity were obtained from the resulting mask, and saved. A calibration (px_movie_ / cm_real object_ ratio) was also obtained from a manually drawn segment and the corresponding length of the object in cm, for later normalization. In following analysis steps, speed and mouse position were derived from the center of gravity coordinates, which were slightly smoothed with a median filter. In addition, to capture general activity, even in the absence of locomotion, a motion measure was used: it is computed as percentage of pixel change in the mouse masks from one frame to the next (non-overlapping pixels / total pixel count). Several body parts (snout, ears, front and hind paws, tail) were also tracked with DeepLabCut^52, 53^. Briefly, a Resnet-152 network was iteratively trained and refined on ∼1350 frames, to be as performant as possible in all our various recording conditions and in particular yield accurate tail tracking (*cf* thermal data extraction). To not sacrifice any accuracy, only coordinates with a score ≥0.99 were included, and no interpolation was applied. A semi-automated threshold-based GUI was used to annotate the following behaviours: rearing, grooming, stretch-attend posture, head dips, immobility, fast locomotion and so-called area-bound (not any of the other defined behaviours, in particular no immobility and no locomotion). Briefly, for each behaviour, a global score was obtained from relevant position information, body parts’ angles / distances / speed, and thresholded with their respective hard-coded thresholds. The behavioural bouts resulting from that initial detection were displayed in a GUI together with the original movie and the scores. The thresholds could then be dragged manually, updating the detected events plots, to get the best possible detection. Events were then checked and adjusted manually when needed within the same GUI, and occasional periods of obstruction (e.g. cable between the camera and the mouse) were marked for later exclusion. In the specific case of the LDB, since the RGB camera wasn’t able to capture mouse’s activity in the dark side, thermal movies were used for behavioural detection. To this end, thermal movies were re-exported with a black-and-white colourmap, after inverting the intensities and adjusting the contrast, so that the resulting frames resemble their RGB counterparts. A DeepLabCut network was derived from our main network, and refined with those new movies (tail points were discarded because of their changing nature on thermal movies). The tracked body parts were then used as previously mentioned to detect behaviours.

#### Spatial analyses

Areas in the different contexts were derived from the contours drawn during tracking, completed if necessary by manual delineation (LDB). In the case of the OF, the center corresponds to the middle square obtained when dividing the OF into a 5 x 5 grid, the corners correspond to the corners of that grid, and the corridors are the squares between two corners. Raw heat maps were generated from the appropriate data and the tracking information using FMA Toolbox (http://fmatoolbox.sourceforge.net/) with 250 bins and no smoothing. A post-processing 2D smoothing was applied, ignoring bins without any occupancy in order to prevent “edge” effects. Corresponding quantification was performed independently from the heat maps, by simply extracting from the raw data the time bins for each area, and processing it accordingly (e.g. averaging). For transition between areas, typical motifs were identified into the list of sequential areas explored by each mouse, and sorted to keep the episodes starting from low speed and without immobility between starting area and target area, except for the OF for which the transition was used as synchronizing event for the PSTH because of the scarcity of events matching the criteria.

### Thermal data extraction

Following acquisition, the large .seq files were read in MATLAB using FLIR SDK (FLIR), and converted to .mj2 files after limiting the values range to 15-50°C and conversion to 8 bits. The thermal movie was subsequently processed using the same GUI as for the RGB top camera, and mouse position, contour, and motion were saved in the same file. Using either synchronization data that was saved online or fitting of the RGB and thermal motion curves, the initial shift and progressive drift between RGB and thermal cameras were determined, and the timestamps adjusted accordingly to align the two cameras temporally. In addition, a Procrustes analysis on the mouse’s global track from both cameras was used to determine the transformation linking the coordinates of both cameras. Checkerboard calibration was performed for later recordings, but Procrustes transform was used over the entirety of the dataset for uniformity. The transformation was then applied to the coordinates of the body parts coming from the RGB camera with DeepLabCut, to obtain their equivalent on the thermal camera, the five points tracked for the tail defining four segments. Although our DeepLabCut network yielded excellent accuracy, and even though the transfer from RGB to thermal camera was reliable, the actual tail width in some large contexts could represent as little as 2 px in the movies. Therefore, the coordinates of a rectangle extending 3 mm on each side of the segment were computed (using the px/mm calibration) for each segment, and the temperature was retrieved as the highest value from the corresponding mask. Because of the 2-D nature of the tracking of a 3D object and of the coordinates transfer, the temperature extraction was subjected to an inherent minimum level of noise (vertical tail, not the same length visible, etc). To correct for it, the data was post-processed to get rid of some artefacts: first, a rank-order filter (90^th^ percentile, 300 samples; Arash Salarian (2021), Rank-Order Filter; https://www.mathworks.com/matlabcentral/fileexchange/22111-rank-order-filter, MATLAB Central File Exchange) was applied to the signal, and a threshold was used to localize artefacts. In a similar manner, moving averages obtained from different window sizes were substracted from the signal, and the results thresholded to identify artefacts. If the resulting ranges were smaller than 15s, missing data was linearly interpolated; otherwise, the values were set to NaN to be discarded from further analyses, and the results were saved.

### Rigidity model and statistics

The term *rigidity* originates from our initial hypothesis that while heart rate can reflect moment-to-moment changes in defensive states, the amplitude does not necessarily reflect only e.g. a discrete “fear level” at a given time, but is also *constrained* by higher-order processes, which was further confirmed and supported by our data: it is a neuro-autonomic phenomenon opposing heart rate changes. An analogy would be a stretched rubber band resisting deflection pressure (Methods Figure 1). Mechanistically, it most likely reflects changes at the level of autonomic balance, which constrains the changes in heart rate that can be achieved –very much like the shifts described for the baroreflex curve^54^. As a result, the action of a ‘microstate generator’, such as the vlPAG neurons responsible for the immobility-bradycardia responses, which always send the same signal (same deflection force onto the rubber band) to its downstream regions, would be strongly modulated by *rigidity* (different pre-load on the rubber band), not always yielding similar amplitude.

**Methods Figure 1.**
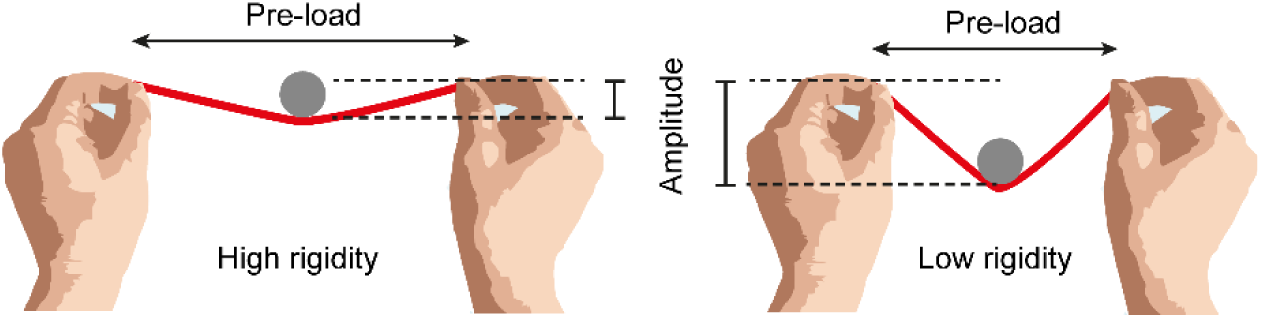
The rubber band analogy for rigidity. When the band is stretched out (high pre-load), it resists deflection force of a weight more and deflection amplitude is low (left). The same weight causes deflection with higher amplitude, when pre-load is lower (right).

Although we show that *rigidity* also works on other microstates (e.g. rearing-associated tachycardia), it is best reflected by immobility-associated bradycardia, a microstate with particular relevance to threat-related reactions. We thus used variations in the amplitude of those cardiac changes as a proxy for *rigidity*.

Because the peak amplitude of immobility-associated bradycardia (*Bradycradia_IB_* measured as beats per minute, bpm) depends both on such within session *rigidity* as well as on bout duration (*Duration_IB_*, we computed coefficients for both factors (SessionCoefficient *A_t_* and BoutDurationCoefficient *B_d_*, respectively). This can be formalized as

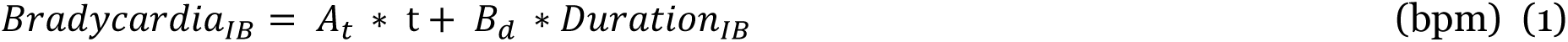

*A_t_* captures the slow decrease in *rigidity,* which can be described as a relaxation process resulting in decreased influence of *rigidity* on bradycardic responses over session time (t). Taking this into account, we introduce *rigidity* as a gain factor, scaling the range (*Bradycradia*_hmax_) of an individual’s possible HR changes. Thus, it constitutes an important parameter defining the momentary ‘state space’ of the cardiac system. In our rubber band analogy, it is the maximal amplitude of deflection. *Rigidity* works against this as the pre-load of the system (or the stretching force on the rubber band), thereby constricting deflection amplitude, or with other words, contracting the state space. Therefore, we define

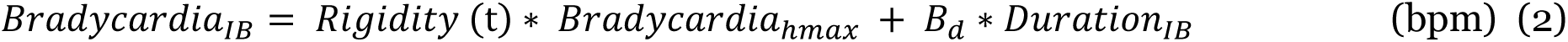

In relation to the SessionCoefficient, this means that

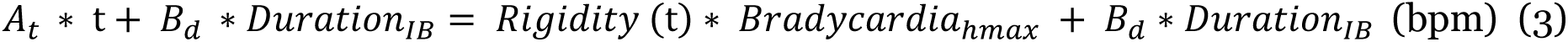

resulting in a dimension-less definition of

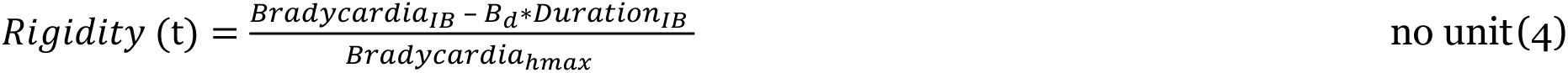

Or

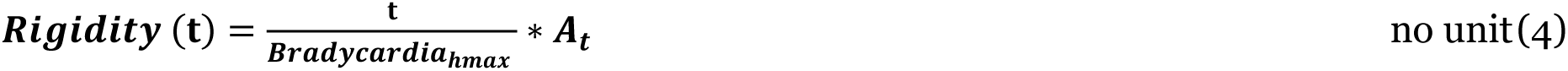

We found immobility-associated bradycardia to depend on both an intrinsic characteristic, namely immobility bout duration, and one general modulatory influence (herein termed *rigidity*). Both to better characterize such defensive response (immobility-associated bradycardia) but also to quantify rigidity, both factors were integrated into a simple model that describes immobility-associated bradycardia’s amplitude in function of time (main contributor to slow rigidity changes) and bout duration.

Practically, to prevent skewing from e.g. individual mice presenting numerous episodes, or single outlier episodes, a fitting wasn’t performed on the pooled individual values per single episode. Instead, average amplitudes of bradycardia were processed for overlapping ranges of immobility bout durations spanning their spread, and at the same time, overlapping time ranges mapping the recording session (e.g. retrieving the minimum value on an average PSTH of HR-to-ceiling for immobility episodes lasting from 2 to 2.5 s and occurring between 0 and 60s would give one data point, with a given bradycardia value for 2.25s bout duration and 30s session time).

The resulting values were fitted using a polynomial equation; robust fitting without normalisation was used (MATLAB Curve fitting toolbox, The MathWorks).

R² values and statistical comparisons where performed on a simple linear model which integrates one constant for each factor (a*BoutDuration + b*SessionTime + c).

We used an F-statistic (Equation 1) to compare different conditions (e.g. open vs closed arm). The fits of the corresponding separate surfaces were compared against the fit of a single surface to fit both conditions (e.g. the whole EPM in this case). A p-value < 0.05 was considered statistically significant, which implicated the datasets were so different that they were best plotted as two separate surfaces.

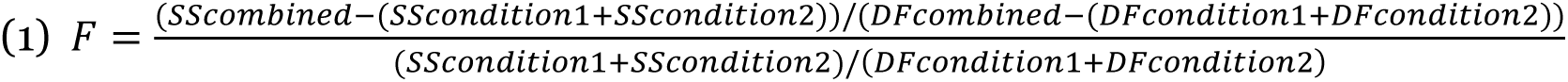

SS, sum of squares error; DF, degrees of freedom.

Adapted from, e.g., Broom, L. *et al.*^55^.

### K-means clustering

The Bayesian Inference Criterion (BIC) was used to determine the optimal number of clusters (Mclust package^56^, in R^57^) on per-mouse averages for conditioning day 2 (speed, motion, HR-to-ceiling, LF amplitude, as well as thermographically recorded tail temperature were used). K-means clustering with the predetermined number of clusters was performed in Matlab (squared Euclidean distance, 1000 replicates).

### UMAP embedding

The different readouts were resampled to a common time vector (4Hz) in MATLAB and exported to a csv file to be embedded with UMAP^39^ (0.5.0) using Euclidean distance, 20 neighbours and minimum distance of 0.3, in either 2 or 3 dimensions. The results of the embedding were exported as xls files and read back into MATLAB were the reduced data was matched with the original readouts and metadata.

For the across-days embedding, 250 UMAP analyses were performed with the relevant data (HR-to-ceiling, immobility bout duration, immobility-associated bradycardia, LF amplitude during locomotion) from all included days each time. The Euclidean distances presented in the matrices are the resulting average of the distances for the 250 runs. Colour-coding for the EPM subareas was added post-hoc by using the metadata.

Embedding for conditioning day 2 was used only for visualization purposes, and performed on speed, motion, HR-to-ceiling, LF amplitude, as well as thermographically recorded tail temperature.

### Specific analyses

For any PSTHs, the appropriate data was extracted around the synchronizing events and resampled to fixed timestamps to allow for cross-trial and cross-animal averaging. 3D representations were built from PSTHs processed over overlapping time windows (60 s width, 50 s overlap).

### Statistics

Normality was checked using Lilliefors test for each set of data, and homoscedasticity was tested with Brown-Forsythe test. When only two sets of data were compared, and if the hypothesis of normality was true, Student’s t test was used, otherwise Wilcoxon signed-rank test or Mann– Whitney U test were used. When more than two sets of data were compared, a one-way ANOVA test was used if the hypotheses of equal variance and normality was true for all, otherwise a Kruskal-Wallis test was used, in both cases followed by appropriate post-hoc test with Bonferroni correction. A mixed model ANOVA was used for the analysis of the behaviours x time interactions supporting the results of Fig. 3a and presented in Extended Data Fig. 3 c,d. For the comparison of the intertrials values as presented in Fig. 6, two-way RM ANOVA followed by Sidak’s multiple comparisons test were performed in Prism 7.05 (GraphPad).

## Supporting information

Heart rate extraction tool

Behavior&HR during conditioning

Optoactivation of Vglut2+ neurons

Optoactivation of Chx10+ neurons

Optoinhibition of Chx10+ neurons

Optoactivation of Vglut2+/Chx10-neurons

## Data availability

Data supporting the findings of this study are available from the corresponding author upon reasonable request.

## Code availability

The custom code used to preprocess ECGs and contour tracking is available at https://github.com/DefenseCircuitsLab/. Other code used in this study is available from the corresponding author upon reasonable request.

## Acknowledgements

We thank K. Walter, E. Bóf-Ramos, H. Troll and C. A. Mehling for technical assistance, and R. Blum for help with microscopy. We thank J. J. Letzkus, K. Heinze, M. S. Esposito, M. S. Fustiñana, J. Müller, M. Schellenberger and C. Redondo Aláñon for input on the manuscript, and all members of the Defense Circuits Lab and Institute of Clinical Neurobiology for discussions and help with the project, in particular M. Sendtner, R. Sendtner and all members of the animal facility. We are grateful for support of this project and JSG by J. Deckert (Center for Mental Health). We also thank the Medical Informatics Service Center (SMI) of the University Hospital Würzburg for providing the infrastructure for data storage and analyses, as well as technical support. We are grateful to O. Kiehn for providing founders of the *Chx10-Cre* mouse line as well as K. Deisseroth and C. Ramakrishnan for sharing the ChRmine AAV and plasmid.

This work was supported by the Deutsche Forschungsgemeinschaft (Heisenberg professorship and project funds to P. Tovote: [TO 1124/1,2,3], TRR 295: [446022135], [446270539]), and received funding from the European Union’s Horizon 2020 research and innovation programme under the Marie Skłodowska-Curie grant agreement No 956414 (P. Tovote). It was supported by a NARSAD Young Investigator Grant of the Brain and Behaviour Foundation to P. Tovote, and a Fellowship grant from the Fundação para a Ciência e a Tecnologia to S.L. Reis.

## Authors information

### Contributions

JSG, NS and PT conceived the project, designed the experiments and wrote the manuscript. NS, SLR and JSG ran behavioural experiments. JSG set up and designed the acquisition and analysis pipeline. DS contributed code for data analysis. JSG, NS, DS and PT analysed the data. SLR, NS and DS performed histochemical analyses.

---

**Extended Data Fig. 1.**
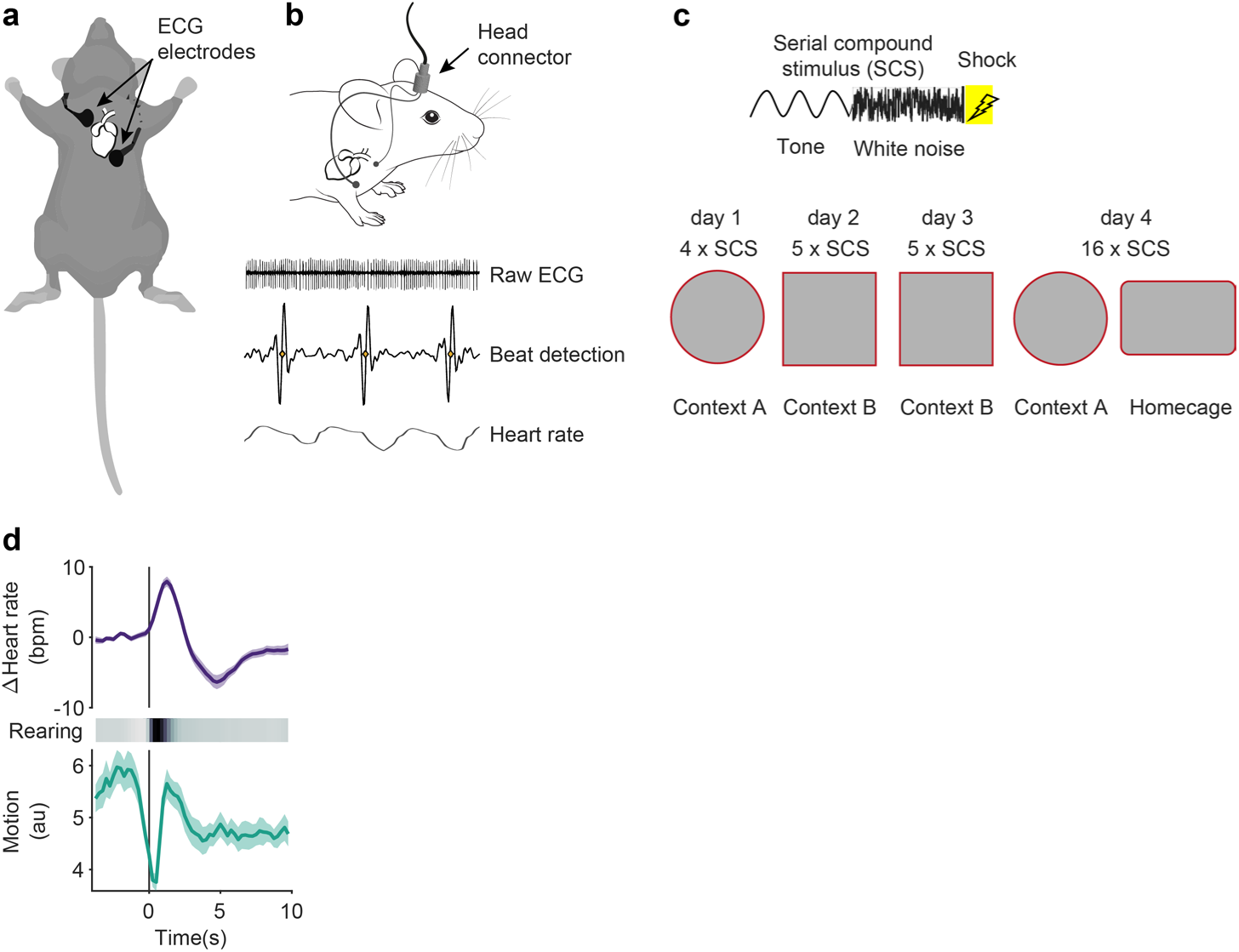
Approaches for cardio-behavioural characterization of defensive states. **a**, Scheme showing the electrodes’ subcutaneous placement for ECG recordings. **b,** Electrodes are soldered to a connector in order to acquire the signal, from which heart beats are extracted to process HR. **c,** Description of the conditioned flight paradigm. Top, the serial compound stimulus consists in pure tone beeps followed by white noise bursts, and shock for conditioning days. Bottom, contexts and number of presentations for the successive days. **d,** PSTHs showing the average HR (top) and behavioural responses (middle and bottom) accompanying rearing bouts (OF, n = 23 mice).

**Extended Data Fig. 2.**
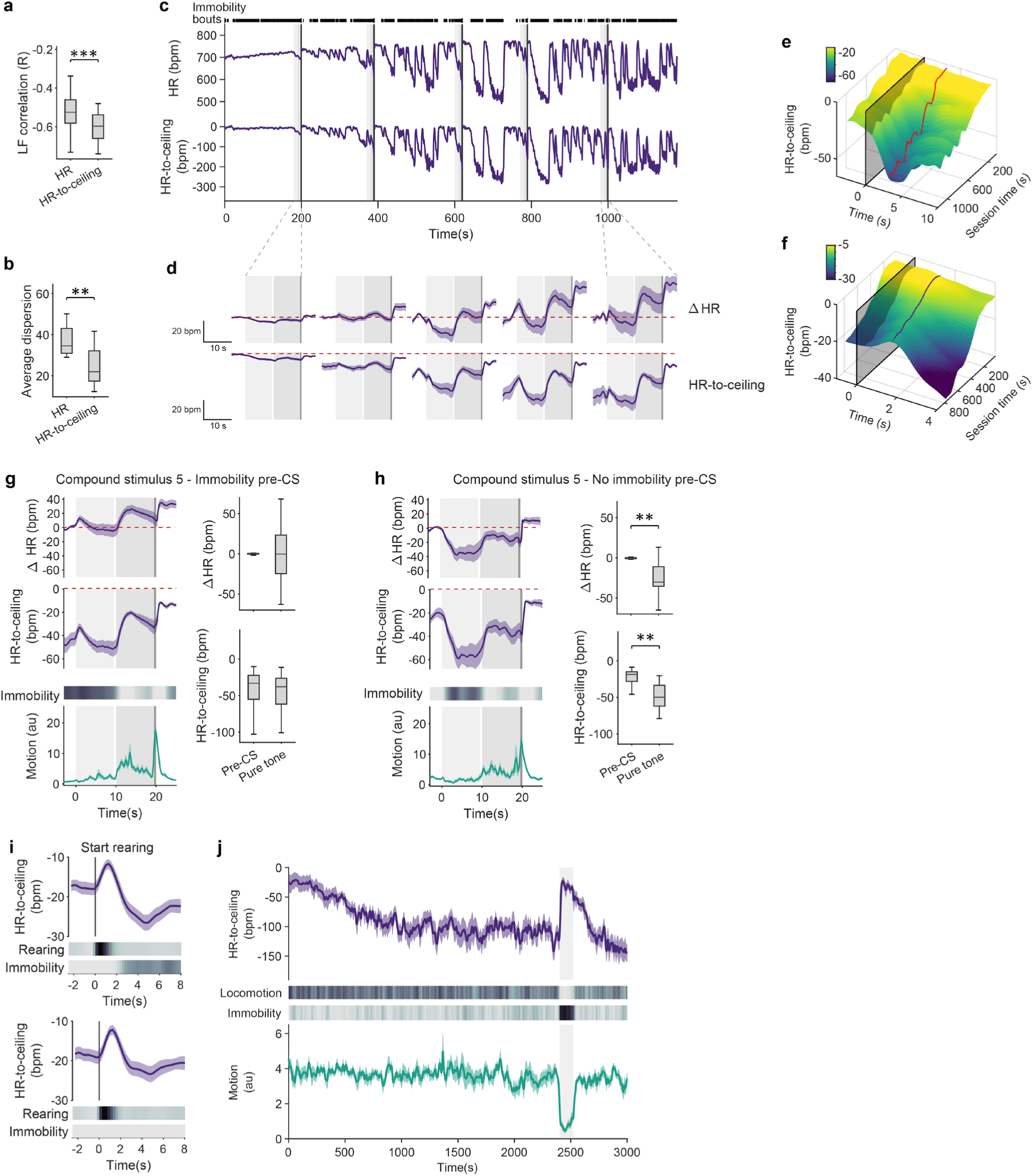
HR-to-ceiling as an alternative to other HR readouts prevents analytical and interpretational confounds, and exposes microstates dependencies. **a-b**, HR-to-ceiling is better correlated to LF amplitude (**a**, n = 33 mice, two-tailed Mann–Whitney test) and explains more variability (**b**, n = 33 mice, two-tailed t-test) than raw HR even when ignoring the initial ramping phase. **c**, Example raw HR (top) and corresponding HR-to-ceiling traces (bottom) from a conditioning day 2 recording for a single mouse; black lines on top represent immobility bouts. **d**, PSTHs of ΔHR (top) or HR-to-ceiling (bottom) for the successive CS-US pairings show a progressive increase in the HR changes’ amplitude but also that ΔHR is misleading by not exposing the initial bradycardia and showing the rest as an increase from baseline, unlike HR-to-ceiling which shows it unambiguously (n = 33 mice). Red dashed line shows pre-SCS1 values to help visualization, and greyed areas represent pure tone, white noise, and shock. **e-f**, 3D representations showing the increase in amplitude of the immobility- and rearing-associated HR changes in function of the recording time as in Fig. 1f and h, respectively, but with HR-to-ceiling instead of ΔHR. HR-to-ceiling also better captures the global relaxation over time as visible in the baseline. **g-h**, PSTHs (left) and matching quantifications (right) for the last CS-US pairings from conditioning day 2 (n = 33 mice), split into trials for which there was immobility before CS onset (g) or not (**h**). Because pre-CS immobility is accompanied by low HR, ΔHR suggests opposite HR changes over the CS-US pairing period (**g** and **h**, top curve) depending on whether there was immobility before or not (g and h, Heat map and bottom curve for motion). HR-to-ceiling (**g** and **h**, middle curve) on the other hand allows to clearly interpret the difference as coming for the pre-CS state of the animal while the CS-US pairing leads to similar changes in both cases (immobility or not before). **i**, PSTHs comparing HR-to-ceiling changes accompanying rearing, whether the bout was closely followed by some immobility (top panel) or not (bottom panel), and showing that immobility accounts for some of the post-rearing bradycardia that can be observed on the overall average. **j**, Average curves for the CS (pure tone in this case) retrieval after a long baseline in the homecage (n = 8 mice). The CS (greyed area) was presented after the HR-to-ceiling had stabilised at a low value (top curve).

**Extended Data Fig. 3.**
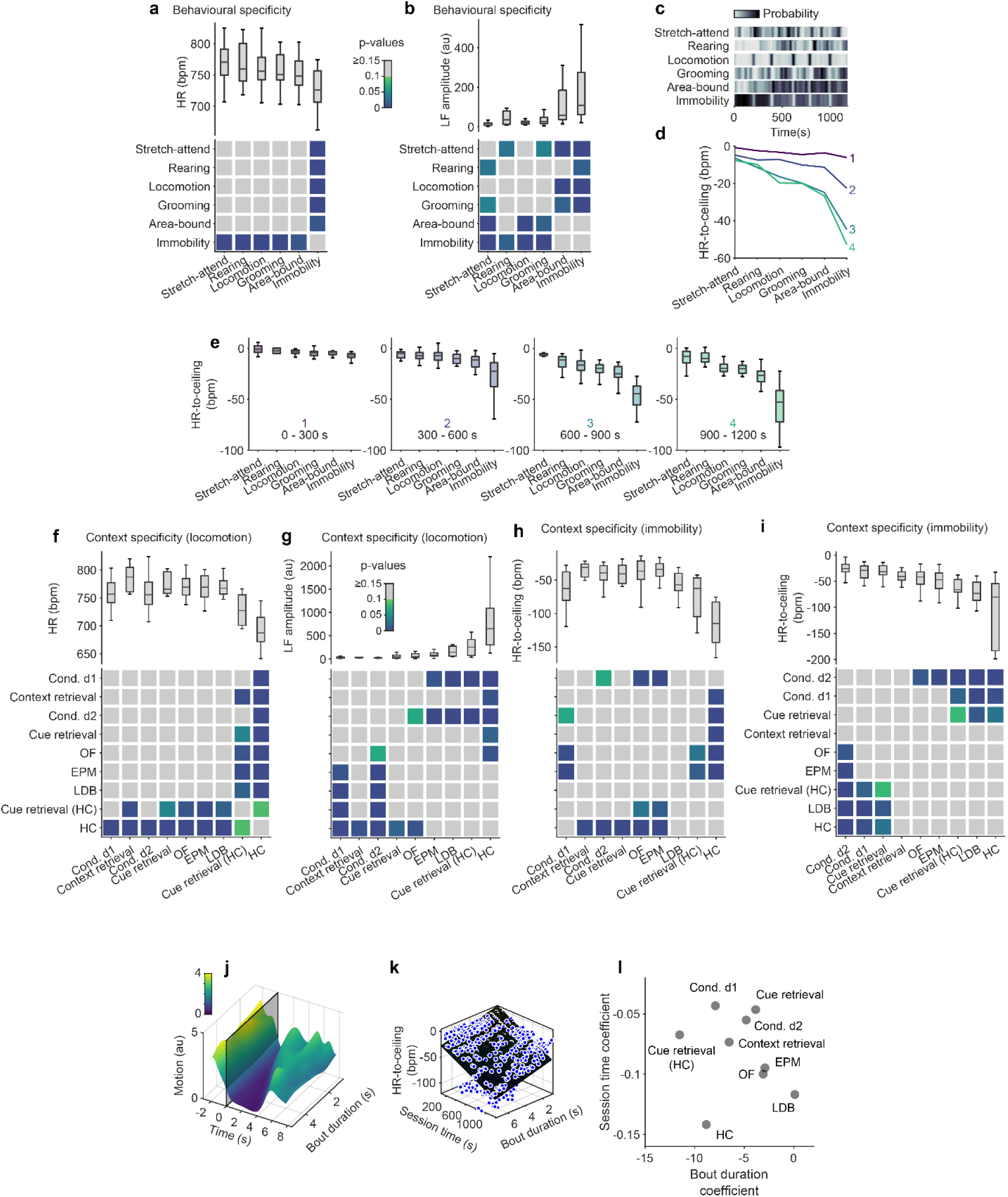
Microstate hierarchies revealed by HR-to-ceiling are recapitulated by LF amplitude but not raw HR, and depend on intrinsic properties of microstates. **a**, Average values of HR for different behaviours during conditioning day 2 recordings (top), and grid showing the results of statistical evaluation of the differences between them (bottom; one-way ANOVA followed by post-hoc pairwise comparison with Bonferroni correction), hinting towards a hierarchy but barely reaching any statistical significance because of high variability. **b**, Average values of LF amplitude for different behaviours during conditioning day 2 recordings (top), and grid showing the results of statistical evaluation of the differences between them (bottom; Kruskal-Wallis followed by post-hoc pairwise comparison with Bonferroni correction). The hierarchy is similar to that of HR-to-ceiling values. **c**, Probabilities of the different behaviours over the course of cond. d2 (n = 33 mice). There are no clear evolution over time that could interact with the progressive rigidity decrease to explain the ranking of the average HR-to-ceiling values (Fig. 3a). **d**, Average values of HR-to-ceiling for the different behaviours. Each curve (1 to 4) represents a different time range within the conditioning day 2 session (respectively 0 - 300 s, 300-600 s, 600-900 s and 900-1200 s). While the values are initially (1, top) constrained to a narrow range by the high rigidity, the differences get more pronounced over time, but the ranking of the different behaviours exhibits stability, demonstrating again that the average in Fig. 3a reflects a global difference for the whole conditioning day 2 (RM ANOVA on n = 33 mice: effect of time and behaviour, p<0.001; interaction, p<0.001) **e**, Similar analyses as Fig.3a, but for the individual time ranges used in (**d**), to better illustrate the ranking of HR-to-ceiling values for the different behaviours is maintained throughout the session. **f**, Comparison of the average values of HR during locomotion in the different paradigms (top), and corresponding statistical analysis (bottom; Kruskal-Wallis followed by post-hoc pairwise comparison with Bonferroni correction). There is a lot of variability and no clear difference nor hierarchy. **g**, Comparison of the average values of LF amplitude during locomotion in the different paradigms (top), and corresponding statistical analysis (bottom; Kruskal-Wallis followed by post-hoc pairwise comparison with Bonferroni correction). The hierarchy parallels that of HR-to-ceiling. **h**, Grand average for HR-to-ceiling values associated with immobility yields paradoxical picture when immobility duration and relative time in the recording are not taken into account (Kruskal-Wallis followed by post-hoc pairwise comparison with Bonferroni correction). i, After accounting for bout duration and relative time during the recordings, HR-to-ceiling values associated with immobility also allow to differentiate the contexts (one-way ANOVA followed by post-hoc pairwise comparison with Bonferroni correction). **j**, 3D representation of motion values in function of immobility bout duration, showing the expected correlation (n = 33 mice). **k**, Same fit as in Fig. 3j, with individual average values displayed as dots. **l**, Session time coefficient in function of bout duration coefficient for each paradigm. Apart from HC-related coefficients which seem to not follow the same logic, bout duration coefficients for the other paradigms follow an opposite ranking as for the session coefficients: higher threat contexts are associated with a larger bout duration coefficient. For **f-I & l**, OF: n= 23 mice; EPM: n = 20 mice; conditioning day 1: n= 30 mice; conditioning day 2: n= 33 mice; LDB: n = 12 mice; context retrieval: n = 5 mice; cue retrieval: n = 10 mice; cue retrieval (HC): n = 9 mice, HC: n = 10 mice.

**Extended Data Fig. 4.**
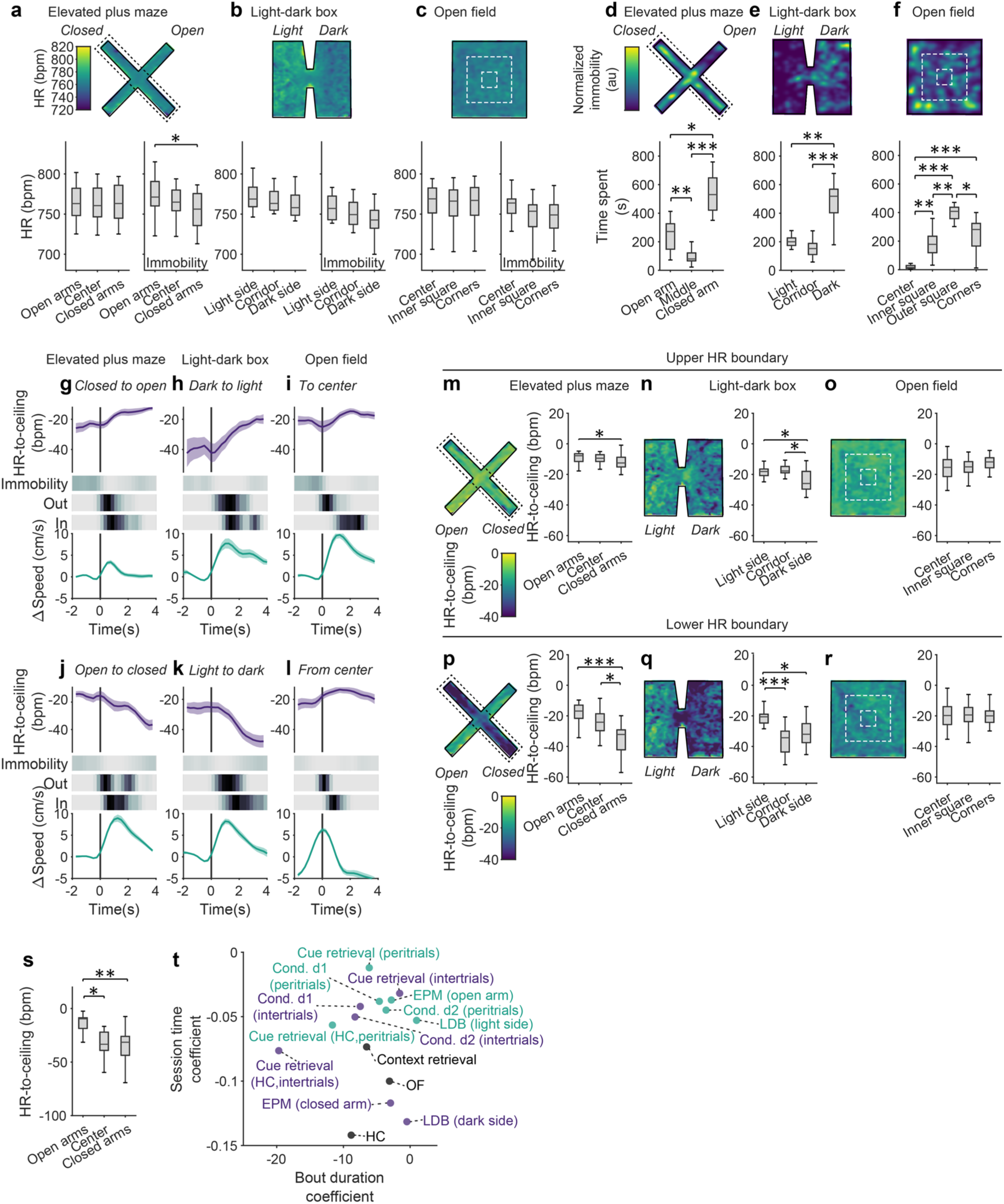
HR-to-ceiling, but not raw HR, reflects subarea-related macrostates, their influence on microstates, and transitions. **a-c**, Heat map representations of the mean HR values in the different contexts (top), corresponding quantification for discrete subareas (bottom, left boxplot; EPM: n = 20 mice, one-way ANOVA; LDB: n = 12 mice, one-way ANOVA; OF: n = 23 mice, one-way ANOVA), and subarea-specific values of HR during immobility (bottom, right boxplot; EPM: n = 20 mice, one-way ANOVA followed by post-hoc pairwise comparison with Bonferroni correction; LDB: n = 12 mice, one-way ANOVA; OF: n = 23 mice, Kruskal-Wallis). **d-f,** Heat map representations of the normalized immobility probability in the different contexts (top), corresponding quantification for discrete subareas (bottom, EPM: n = 20 mice, one-way ANOVA; LDB: n = 12 mice, one-way ANOVA; OF: n = 23 mice, one-way ANOVA). **g-l**, PSTHs showing separately average HR-to-ceiling (top) and speed changes (bottom) for the transitions from low threat areas to higher threat areas (top row) or the opposite (bottom row), for EPM (n = 20 mice), LDB (n = 12 mice) and OF (n = 23 mice). The heat plots below immobility probability show the cumulative distribution for all the averaged transitions of the exit from the start area (“Out”) and the arrival in the target area (“In”). Note that both modest immobility probability, and conversely, the ongoing locomotion, are similar for both directions, or that higher speed is associated with the bradycardic response, and that locomotion is therefore unlikely to account for the changes in HR-to-ceiling observed. **m-r**, Heat map representations of the maximum (top row) as well as minimum (bottom row) HR-to-ceiling values in the different contexts, and corresponding quantification for discrete subareas (EPM: n = 20 mice, Kruskal-Wallis (top) and one-way ANOVA (bottom) followed by post-hoc pairwise comparison with Bonferroni correction; LDB: n = 12 mice, Kruskal-Wallis followed by post-hoc pairwise comparison with Bonferroni correction; OF: n = 23 mice, one-way ANOVA). **s,** HR-to-ceiling average values associated with immobility episodes in subareas of the EPM (n = 20 mice, Kruskal-Wallis followed by post-hoc pairwise comparison with Bonferroni correction); similar to Fig. 4j, except that only immobility episodes of similar duration were selected here, to confirm that the effect observed is not due to a potential difference in bout duration between subareas. **t,** Session time coefficient in function of bout duration coefficient for each paradigm and/or subperiod/subarea. While the ranking for session coefficient follows extremely well the intuitive threat levels, bout duration coefficient is likely to be influenced by other factors that are not readily visible in this analysis.

**Extended Data Fig. 5.**
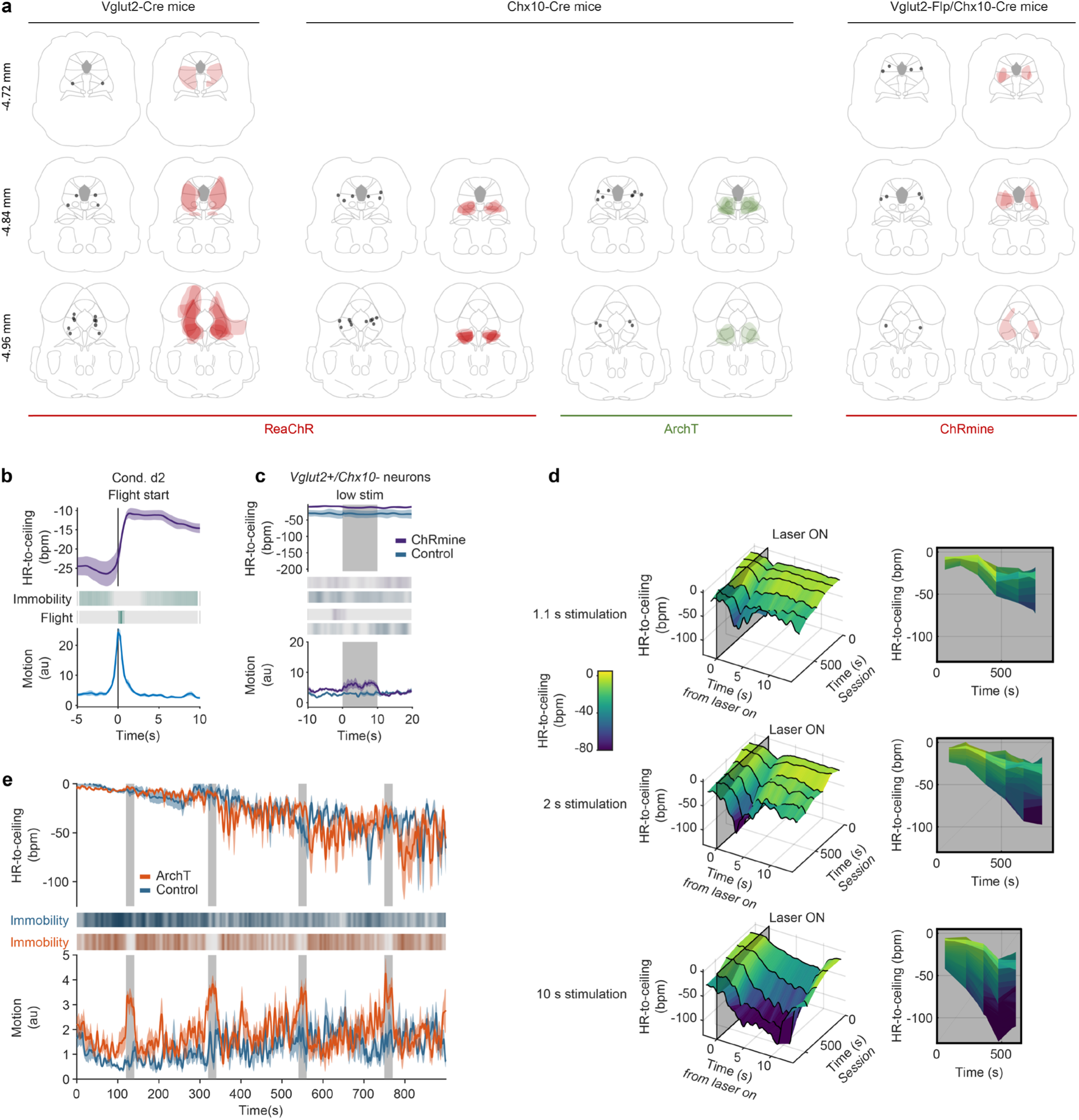
Histology for the optogenetics experiment and rigidity effect generalization to different stimulation protocols. **a**, For each combination of mouse line and viral approach used for the optogenetics experiments, one line per brain level, where the left midbrain section shows the fiber placement and the right one depicts viral expression as overlay for all the mice. **b**, PSTHs showing the average HR (top) and behavioural responses (middle and bottom) associate with flight during conditioning day 2 (n = 33 mice). **c**, PSTHs depicting the responses evoked by a low intensity stimulation of ChRmine-expressing *Vglut2+/Chx10-* neurons in the vlPAG. Only a mild behavioural activation was visible on the average motion curve (bottom). **d**, 3D representations of the amplitude of HR decrease evoked by optogenetic stimulation of the *Chx10+* neurons (n = 8 mice) in function of the relative time from the beginning of the recording (left panels), and corresponding 2D views (right panels), for the 1.1, 2 and 10 s stimulations. The amplitude of bradycardia increases with time, revealing it is affected by rigidity even for more modest stimulation durations, in a linear manner. **e,** Average traces for the whole recordings for the optogenetic inhibition of the *Chx10+* neurons from the vlPAG during context retrieval, showing the inhibition of immobility by the stimulations (grey areas).

**Extended Data Fig. 6.**
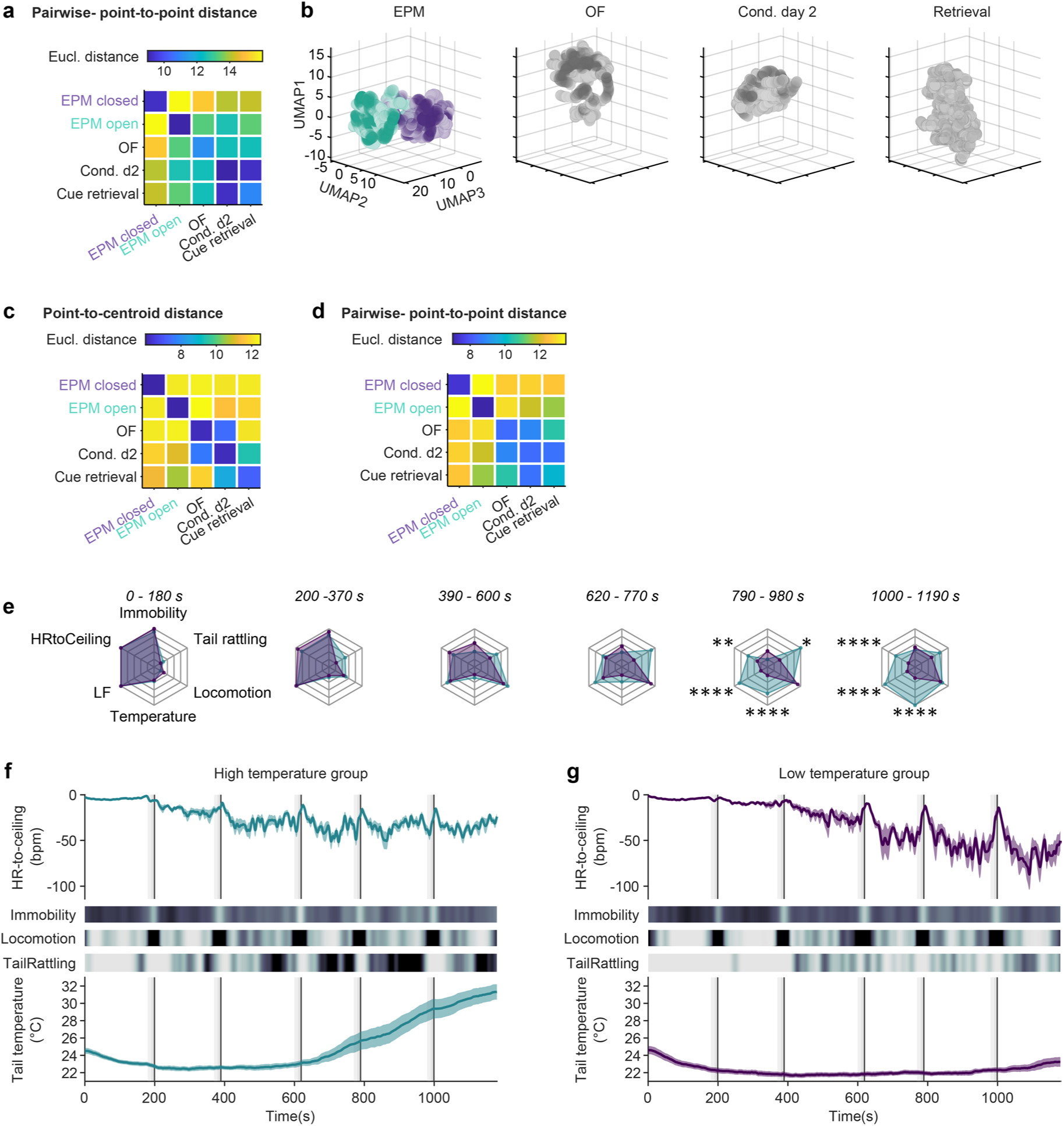
Extension to different metrics and dimensionalities of the “state space”-defining methodology and detailed representation of the raw data from the diverging subgroups. **a**, Matrix showing the average pairwise Euclidean distance between single data points for the different contexts corresponding to the UMAP results in Fig. 6a (average of 250 UMAP runs). **b**, 3D representation of the UMAP dimensionality reduction of secondary readouts of interest (ceiling, immobility bout duration, immobility-associated bradycardia, LF amplitude during locomotion) extracted for different paradigms (similar to Fig. 6a, but new analysis with a reduction to three dimensions instead of two). The two-coloured sub-clouds for the EPM correspond to the open arm and closed arm data points. c-d, Matrix showing the average Euclidean distance between each contexts’ data points and the others’ centroids (**c**) or average pairwise Euclidean distances between single data points (**d**), corresponding to the UMAP results in **b** (averages of 250 UMAP runs). **e**, Spider charts showing the average values of several readouts over the 6 ISI for the two groups of mice identified (respectively n = 14 vs n = 15), with progressively pronounced autonomic and behavioural differences (two-way RM ANOVA followed by Sidak’s multiple comparisons test). **f-g**, Average curves of HR-to-ceiling (top), immobility, locomotion, and tail rattling probabilities (heat plots in the middle) and tail temperature (bottom) for the two groups of mice, one of which presented an increase in tail temperature (**f**, n = 14) while the other did not (**g**, n = 15).

**Extended data Table 1.**

| Figure | Normality test | Pairwise test | Comparison | Tail | p-value | Group analysis | F-statistic (or Chi-square) | p-value |
| --- | --- | --- | --- | --- | --- | --- | --- | --- |
| 2b | passed | unpaired Student's t-test | HR vs HR-to-ceiling | two-tailed | 1.2291e-07 |  |  |  |
| 3a | failed | post-hoc with Bonferroni correction | Behaviours | two-tailed | cf matrix | Kruskal-Wallis | (5,164) = 118.7625 | 5.7388e-24 |
| 3b | failed | post-hoc with Bonferroni correction | Contexts | two-tailed | cf matrix | Kruskal-Wallis | F(8,145) = 81.8844 | 2.0398e-14 |
| 4a | passed | post-hoc with Bonferroni correction | EPM subareas | two-tailed | Open arm vs closed arm: 5.57469e-05<br>Center vs closed arm: 0.02329 | one-way ANOVA | F(2, 60) = 10.9428 | 8.8782e-05 |
| 4b | passed | post-hoc with Bonferroni correction | LDB subareas | two-tailed | Light vs corridor: 0.033799<br>Light vs dark: 0.0092517 | one-way ANOVA | F(2, 42) = 5.7107 | 0.0064004 |
| 4c | passed | - | OF subareas | - | - | one-way ANOVA | F(2, 66) = 0.21763 | 0.805 |
| 4j | passed | post-hoc with Bonferroni correction | EPM subareas | two-tailed | Open arm vs closed arm: 2.1636e-05<br>Center vs closed arm: 0.012626 | one-way ANOVA | F(2, 56) = 12.377 | 3.5372e-05 |
| 4k | passed | - | LDB subareas | - | - | one-way ANOVA | F(2, 30) = 2.3881 | 0.10904 |
| 4l | passed | - | OF subareas | - | - | one-way ANOVA | F(2, 58) = 0.23383 | 0.79224 |
| 4d top | passed | unpaired Student's t-test | Closed to open vs open to closed | two-tailed | 1.3966e-10 |  |  |  |
| 4d bottom | failed | Mann-Whitney | Closed to open vs open to closed | two-tailed | 3.9446e-07 |  |  |  |
| 4e top | passed | unpaired Student's t-test | Light to dark vs dark to light | two-tailed | 1.6518e-04 |  |  |  |
| 4e bottom | passed | unpaired Student's t-test | Light to dark vs dark to light | two-tailed | 0.7214 |  |  |  |
| 4f top | failed | Mann-Whitney | Inner to outer vs outer to inner | two-tailed | 0.3345 |  |  |  |
| 4f bottom | failed | Mann-Whitney | Inner to outer vs outer to inner | two-tailed | 0.0022586 |  |  |  |
| 4g | passed (no homoscedasticity) | post-hoc with Bonferroni correction | EPM subareas | two-tailed | Open arm vs center: 0.0995780<br>Open arm vs closed arm: 4.1001e-07 | Kruskal-Wallis | (2,60) = 28.1091 | 7.8737e-07 |
| 4h | passed (no homoscedasticity) | post-hoc with Bonferroni correction | LDB subareas | two-tailed | Light vs corridor: 2.3937e-08<br>Light vs dark: 0.0084048<br>Dark vs corridor: 0.016298 | Kruskal-Wallis | (2,42) = 33.2947 | 5.8905e-08 |
| 4i | failed | post-hoc with Bonferroni correction | OF subareas | two-tailed | Center vs corners: 2.8572e-04<br>Center vs inner ring: 0.0011572 | Kruskal-Wallis | (2,66) = 18.6355 | 8.9816e-05 |
| 5g | passed | unpaired Student's t-test | Controls vs ArchT | two-tailed | 0.0027 |  |  |  |
| Ext. Data 2a | failed | Mann-Whitney | HR vs HR-to-ceiling | two-tailed | 8.8985e-04 |  |  |  |
| Ext. Data 2b | passed | unpaired Student's t-test | HR vs HR-to-ceiling | two-tailed | 0.0017082 |  |  |  |
| Ext. Data 2g top | passed | paired Student's t-test | Pre-CS vs pure tone | two-tailed | 0.86001 |  |  |  |
| Ext. Data 2h top | passed | paired Student's t-test | Pre-CS vs pure tone | two-tailed | 0.0042966 |  |  |  |
| Ext. Data 2g bottom | failed | Wilcoxon signed-rank | Pre-CS vs pure tone | two-tailed | 0.81408 |  |  |  |
| Ext. Data 2h bottom | passed | paired Student's t-test | Pre-CS vs pure tone | two-tailed | 0.0041187 |  |  |  |
| Ext. Data 3a | passed | post-hoc with Bonferroni correction | Behaviours | two-tailed | cf matrix | one-way ANOVA | $F(5,164) = 8.2619$ | 5.5314e-07 |
| Ext. Data 3b | failed | post-hoc with Bonferroni correction | Behaviours | two-tailed | cf matrix | Kruskal-Wallis | $(5,164) = 74.1437$ | 1.4037e-14 |
| Ext. Data 3d | - | post-hoc with Tukey's test | Behaviours, time | two-tailed | - | RMANOVA (mixed model) | Behaviours: $F(1.661, 202.1) = 58.43$<br>Time: $F(5, 165) = 42.95$<br>Interaction: $F(15, 365) = 14.08$ | All <0.0001 |
| Ext. Data 3f | failed | post-hoc with Bonferroni correction | Contexts | two-tailed | cf matrix | Kruskal-Wallis | $(8, 145) = 77.0866$ | 7.5145e-13 |
| Ext. Data 3g | failed | post-hoc with Bonferroni correction | Contexts | two-tailed | cf matrix | Kruskal-Wallis | $(8, 145) = 77.0866$ | 7.5145e-13 |
| Ext. Data 3h | failed | post-hoc with Bonferroni correction | Contexts | two-tailed | cf matrix | Kruskal-Wallis | $(8, 145) = 57.2016$ | 1.6447e-09 |
| Ext. Data 3i | passed (no homoscedasticity) | post-hoc with Bonferroni correction | Contexts | two-tailed | cf matrix | Kruskal-Wallis | $F(8,145) = 60.9944$ | 2.9733e-10 |
| Ext. Data 4a left | passed | - | EPM subareas | - | - | one-way ANOVA | $F(2, 60) = 0.048052$ | 0.95312 |
| Ext. Data 4a right | passed | post-hoc with Bonferroni correction | EPM subareas | two-tailed | Open arm vs closed arm: 0.02808 | one-way ANOVA | $F(2, 55) = 3.632$ | 0.032997 |
| Ext. Data 4b left | passed | - | LDB subareas | - | - | one-way ANOVA | $F(2, 42) = 0.7631$ | 0.47257 |
| Ext. Data 4b right | passed | - | LDB subareas | - | - | one-way ANOVA | $F(2, 30) = 2.0124$ | 0.15131 |
| Ext. Data 4c left | passed | - | OF subareas | - | - | one-way ANOVA | $F(2, 66) = 0.022819$ | 0.97745 |
| Ext. Data 4c right | failed | - | OF subareas | - | - | Kruskal-Wallis | $(2, 58) = 4.1024$ | 0.12858 |
| Ext. Data 4d bottom | failed | post-hoc with Bonferroni correction | EPM subareas | two-tailed | Open arm vs middle: 0.0046<br>Open arm vs closed arm: 0.0134<br>Closed arm vs middle: <0.001 | Kruskal-Wallis | $F(2, 54) = 36.1628$ | 1.404e-08 |
| Ext. Data 4e bottom | failed | post-hoc with Bonferroni correction | LDB subareas | two-tailed | Light vs dark: 0.0017<br>Corridor vs dark: <0.001 | Kruskal-Wallis | $F(2, 51) = 32.3964$ | 9.2302e-08 |
| Ext. Data 4f bottom | failed | post-hoc with Bonferroni correction | OF subareas | two-tailed | Center vs inner square: 0.0011<br>Center vs outer square: <0.001<br>Center vs corners: <0.001<br>Inner square vs outer square: 0.0046<br>Outer square vs corners: 0.0329 | Kruskal-Wallis | $F(3,72) = 51.1569$ | 4.5296e-11 |
| Ext. Data 4m | failed | post-hoc with Bonferroni correction | EPM subareas | two-tailed | Open arm vs closed arm: 0.022858 | Kruskal-Wallis | $(2, 60) = 7.5139$ | 0.023355 |
| Ext. Data 4n | passed<br>(no homoscedasticity) | post-hoc with Bonferroni correction | LDB subareas | two-tailed | Corridor vs dark:<br>0.034221s | Kruskal-Wallis | (2, 42) = 7.6274 | 0.022066 |
| Ext. Data 4o | passed | - | OF subareas | - | - | one-way ANOVA | F(2, 66) = 1.56 | 0.21779 |
| Ext. Data 4p | passed | post-hoc with Bonferroni correction | EPM subareas | two-tailed | Open arm vs closed arm:<br>4.1732e-06<br>Center vs closed arm:<br>0.014911 | one-way ANOVA | F(2, 60) = 14.4068 | 7.7652e-06 |
| Ext. Data 4q | passed<br>(no homoscedasticity) | post-hoc with Bonferroni correction | LDB subareas | two-tailed | Light vs corridor:<br>0.00033399<br>Light vs dark: 0.014955 | Kruskal-Wallis | (2, 42) = 15.9567 | 0.0003428 |
| Ext. Data 4r | passed | - | OF subareas | - | - | one-way ANOVA | F(2, 66) = 0.13698 | 0.87223 |
| Ext. Data 4s | failed | post-hoc with Bonferroni correction | EPM subareas | two-tailed | Open arm vs center:<br>0.023973<br>Open arm vs closed arm:<br>0.0027776 | Kruskal-Wallis | (2,34) = 12.51 | 0.0019208 |

